# Clec7a-mediated regulation of Killer-like Lectin Receptor expression controls T cell immunity

**DOI:** 10.64898/2025.12.04.692469

**Authors:** Ivy M. Dambuza, Fabián Salazar, Emily A. Sey, Stavrola L. Kastora, Cecilia Rodrigues, Annie Phillips-Brookes, Jamie Harvey, Rebecca A. Drummond, Jane Rush, Raif Yuecel, Shinobu Saijo, Janet A. Willment, Daniel H. Kaplan, Salomé LeidundGut-Landmann, Gordon D. Brown

## Abstract

Clec7a is a C-type lectin receptor (CLR) originally defined for its non-redundant role in anti-fungal immunity. Subsequent work has broadened this view, implicating Clec7a in host defense against diverse pathogens and in the pathogenesis of cancer, autoimmunity, neuroinflammation, and developmental disorders. How a single innate receptor orchestrates such wide-ranging outcomes remains unresolved. We previously demonstrated that dendritic cell (DC)-expressed Clec7a is required for protective anti-fungal immunity in the gastrointestinal tract through regulation of fungus-specific CD4⁺ T cell responses. Here, we show that Clec7a controls the expression of multiple C-type lectins in DCs, including a cluster of killer lectin-like receptors (KLRs). Notably, we reveal that these KLRs directly regulate DC function and control CD4⁺ T cell responses. These findings define a novel Clec7a-KLR axis that integrates innate and adaptive immunity, highlighting a regulatory pathway with broad relevance for immune homeostasis, inflammation, and host defense.

## Main

The orchestration of adaptive immunity depends on dendritic cells (DCs) integrating pathogen-derived signals into tailored instructions for T cell activation and differentiation. Among DC pattern recognition receptors, Clec7a (Dectin-1) is a prototypic C-type lectin receptor that recognizes fungal β-glucans and was initially defined by its role in innate responses, including phagocytosis, reactive oxygen species production, and pro-inflammatory cytokine secretion through Syk-CARD9 and Raf-1 pathways^1, 2, 3, 4^. Beyond innate immunity, Clec7a is now firmly established as a regulator of adaptive T cell responses. In DCs, its signalling promotes cytokine programs that drive Th1 and Th17 polarization^5, 6, 7, 8^. Human genetics underscores this non-redundant role of Dectin-1 in immunity against fungal infection as individuals carrying *CLEC7A* loss-of-function mutations (e.g., Y238X) exhibit defective cytokine production including IL-6, IL-17, IL-1β and TNF production by peripheral blood mononuclear cells (PBMCs) and macrophages and are highly susceptible to fungal infections^9, 10^. Together with other CLRs, such as Mincle, Clec7a can also drive development of Th2 responses^11^. Intriguingly, in mouse models of fungal infection, lack of Clec7a has also been linked with dysregulated elevation of Th1 immunity, IFN-γ and IL-17A^12, 13^.

Notably, the influence of Clec7a extends beyond anti-fungal immunity. Clec7a has been implicated in host responses to bacteria, viruses, and parasites, as well as in non-infectious pathologies including autoimmunity, asthma, neuroinflammation, and cancer^14^. Despite these diverse associations, the mechanisms by which Clec7a controls such broad immune outcomes remains poorly defined. Most studies have focused on Clec7a’s role in inducing cytokines like IL-6, IL-12, and IL-23, which tailor CD4^+^ T cell responses. However, these alone cannot fully explain the spectrum of CD4^+^ T cell responses attributed to Clec7a. Our work previously demonstrated that DC-expressed Clec7a is required for maintaining protective anti-fungal immunity in the gastrointestinal tract by promoting survival of fungus-specific CD4^+^ T cells^15^. Here, we wanted to understand how a single innate receptor is acting as a regulator of CD4^+^ T cell immunity, as dysregulated CD4^+^ T cell responses underlie not only susceptibility to fungal pathogens but also the immunopathology of autoimmunity, chronic inflammation, and cancer. We have discovered that Clec7a controls the expression of multiple genes in DCs, including a cluster of CLRs known as killer lectin-like receptors (KLRs). We show a previously unrecognized Clec7a-KLR axis that directly regulates CD4^+^ T cell activation, differentiation, and survival.

## Results

### Clec7a regulates CD4^+^ T cell activation

Using the OT-II system and a *Candida albicans* strain expressing ovalbumin, we previously described how Clec7a deficiency results in the loss of antigen-specific CD4^+^ T cells in the gastrointestinal tract (GIT) and mesenteric lymph nodes (mLN) during systemic infection^15^. To further investigate the underlying mechanisms, we employed the *C. albicans*-specific Hector TCR transgenic mouse, which expresses a TCR that recognizes *C. albicans* alcohol dehydrogenase 1 (ADH1)^16^. We adoptively transferred naïve 1×10^6^ Hector CD4^+^ T cells into WT and Clec7a^-/-^ mice, and then intravenously infected the mice with 2-2.5×10^5^ *C. albicans* SC5314 yeasts (**Fig. 1a**). As before^15^, we found significantly higher fungal burdens in the intestine of Clec7a^-/-^ mice compared to WT mice accompanied by loss in intestinal tissue mass (**Supplementary Fig. 1a**). Additionally, we observed significantly increased counts of apoptotic Hector CD4^+^ T cells in the mLN of Clec7a^-/-^ mice (**Supplementary Fig. 1b**), confirming our earlier findings using the OT-II system and OVA-*C. albicans*^15^.

**Fig 1.**
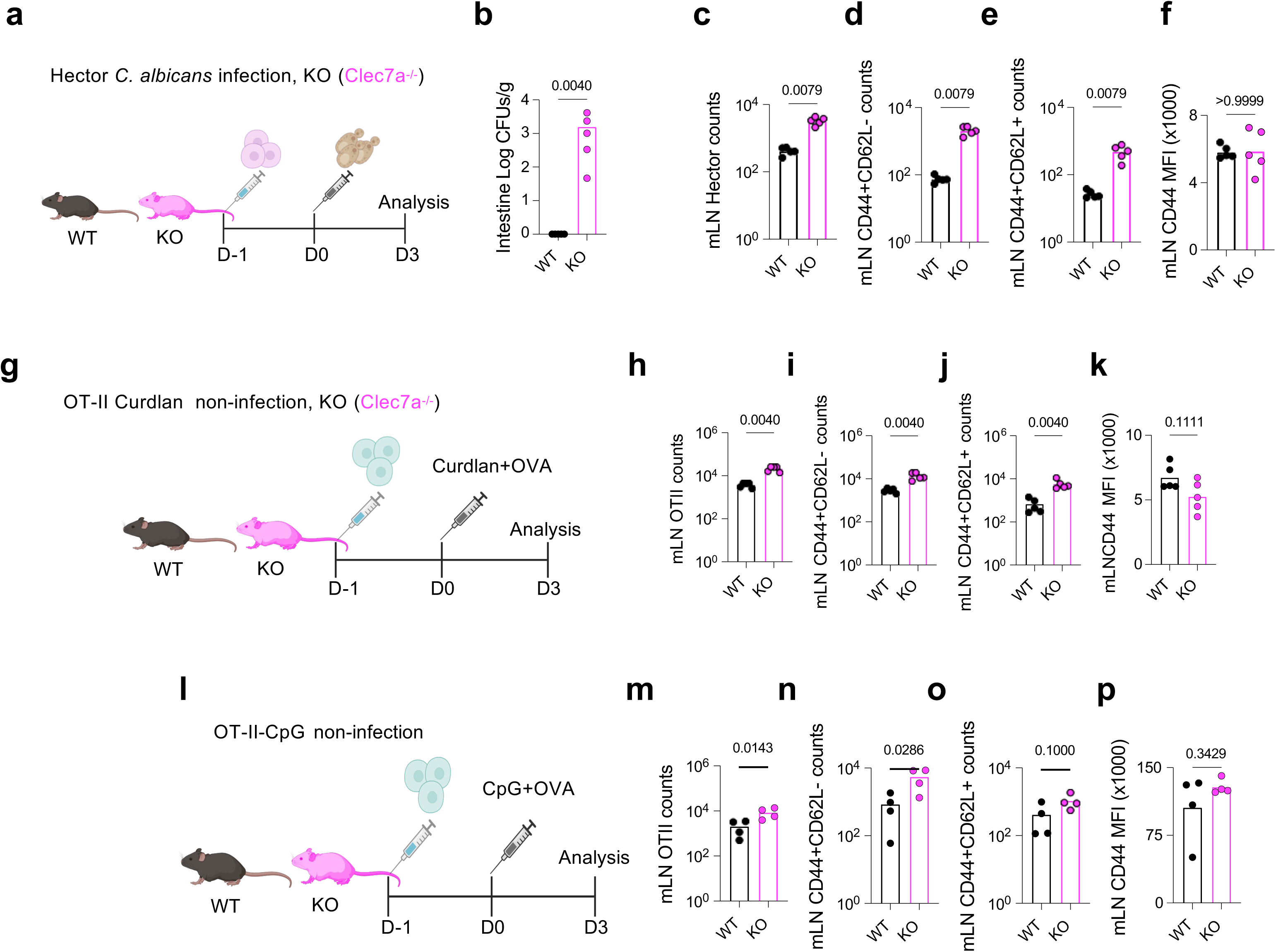
Clec7a regulates CD4^+^ T cell activation. (**a**) Experimental schematic: adoptive transfer of 1×10^6^ naïve Hector CD4^+^ T cells intravenously (i.v) into WT mice and Clec7a^-/-^mice at day-1, and next day i.v infection with 1×10^5^ *C. albicans* SC5314 yeasts. (**b**) Intestine fungal burden 3 days post infection (p.i). Mesenteric lymph nodes (mLN) flow cytometric analysis of number of live (**c**) *C. albicans*-specific Hector CD4⁺ T cells; (**d**) effector memory CD44^+^CD62L^-^; (**e**) central memory CD44^+^CD62L^+^ and (**f**) CD44 mean fluorescence intensity (MFI) analysed within the CD44^+^CD62L^-^ subset of Hector cells. (**g**) OTII-Curdlan challenge schematic: adoptive transfer of 1×10^6^ naïve OT-II CD4^+^ T cells i.v into WT mice and Clec7a^-/-^mice at day-1, and next day intraperitoneal (i.p) challenge with 2 mg/ml curdlan plus 50 ug OVA in 100 ul of phosphate buffered saline (PBS). mLN flow cytometric analysis of number of live (**h**) OVA*-*specific OT-II CD4⁺ T cells; (**i**) effector memory CD44^+^CD62L^-^; (**j**) central memory CD44^+^CD62L^+^ and (**k**) CD44 MFI analysed within the CD44^+^CD62L^-^ subset of OT-II cells. (**l**) OTII-CpG challenge schematic: adoptive transfer of 1×10^6^ naïve OT-II CD4^+^ T cells i.v into WT mice and Clec7a^-/-^mice at day-1, and next day i.p challenge with 50 ug/ml CpG ODN 2395 plus 50 ug OVA in 100 ul of PBS. mLN flow cytometric analysis of number of live (**m**) OVA*-*specific OT-II CD4⁺ T cells; (**n**) effector memory CD44^+^CD62L^-^; (**o**) central memory CD44^+^CD62L^+^ and (**p**) CD44 MFI analysed within the CD44^+^CD62L^-^ subset of OT-II cells. In all the models, analyses were performed on day 3 post challenge with *C. albicans*, or OVA-curdlan, or OVA-CpG. Each point represents one mouse, and horizontal bars indicate the mean. Data are representative of one of three independent experiments. Statistical analysis was performed using Student’s *t*-test with p ≤ 0.05 considered significant.

To explore the possibility that loss of antigen-specific CD4^+^ T cells in the mLN was being driven by high fungal burdens in the GIT, we then challenged mice with a lower inoculum of 1×10^5^ *C. albicans* yeasts. While fungal burdens remained significantly elevated in Clec7a^-/-^ mice (**Fig. 1b**), we observed a significant increase in live *C. albicans*-specific Hector CD4^+^ T cells in the mLN of Clec7a^-/-^ mice compared to WT mice (**Fig. 1c)**. Additionally, in mLN of Clec7a^-/-^ mice, Hector cells displayed enhanced activation with significant numbers of effector memory CD44^+^CD62L^-^ and central memory CD44^+^CD62L^+^ phenotype when compared to Hector cells in WT mice following *C. albicans* infection (**Fig. 1d-e, Supplementary Fig 1c**). Notably, the surface expression of CD44, assessed by mean fluorescence intensity (MFI) within the CD44⁺CD62L⁻ subset of Hector cells, was comparable between Clec7a^-/-^ and WT mice in mLN (**Fig. 1f**). These data suggest that the influence of Clec7a on T cell activation occurs downstream or independently of CD44 modulation. We made similar observations in the spleens of infected Clec7a^-/-^ mice compared to WT mice (**Supplementary Fig. 1d-f**). Furthermore, we found increased numbers of effector memory CD44^+^CD62L^-^ T cells following low-dose infection in the OT-II OVA-*C. albicans* system in the mLN and spleens of Clec7a^-/-^ mice compared to WT mice (**Supplementary Fig. 1g-i**). Together, these confirm that in the absence of Clec7a, the expansion of antigen-specific effector memory CD44^+^CD62L^-^ is not restricted to a specific TCR transgenic model.

To determine whether infection was required for the observed CD4^+^ T cell response, we next employed a non-infectious fungal stimulus. Mice were intraperitoneally challenged with β-glucan curdlan, a Clec7a agonist, to assess T cell activation in the absence of live infection (**Fig. 1g**). Phenotypic analysis revealed that curdlan-challenged Clec7a^-/-^ mice displayed significantly increased numbers of antigen-specific OT-II cells (**Fig. 1h**), effector memory CD44^+^CD62L^-^ cells (**Fig. 1i**), central memory CD44^+^CD62L^+^ cells (**Fig. 1j**) with no change in CD44 MFI expression (**Fig. 1k**), comparable to effects observed during *C. albicans* infection. Next, we employed a DC activating stimulus that does not bind Clec7a by using intraperitoneal administration of CpG^5^ (**Fig. 1l**). As with the *C. albicans* infection or curdlan model, detailed above, we observed increased numbers of activated antigen-specific OT-II cells in the mLN of Clec7a^-/-^ mice compared to WT mice (**Fig. 1m**), effector memory CD44^+^CD62L^-^ cells (**Fig. 1n**), and a trend towards increased central memory CD44^+^CD62L^+^ cells (**Fig. 1o**). There was no change in CD44 MFI expression (**Fig. 1p**). Together, these results reveal that loss of Clec7a is linked with aberrant amplification of antigen-specific CD4^+^ T cell response, irrespective of the innate trigger.

### Altered expression of Killer Like Receptors (KLRs) in Clec7a deficient mice

We previously demonstrated that the altered CD4^+^ T cell response observed in Clec7a^-/-^ mice stems from the DC compartment^15^. To investigate how Clec7a^-/-^ DCs might influence T cell responses, we evaluated their ability to activate naïve (CD44^-^CD62L^+^) CD4^+^ T cells *in vitro* (**Fig. 2a**). Strikingly, CD11c^+^ DCs isolated from the mLN of naïve Clec7a^-/-^ mice induced significantly greater T cell activation compared to WT DCs (**Fig. 2b**), reflecting our *in vivo* observations. To understand the mechanism by which absence of Clec7a mediates increased T cell activation, we assessed the expression levels of the common members of the B7 and TNFR co-signalling molecules on DCs.

**Fig. 2.**
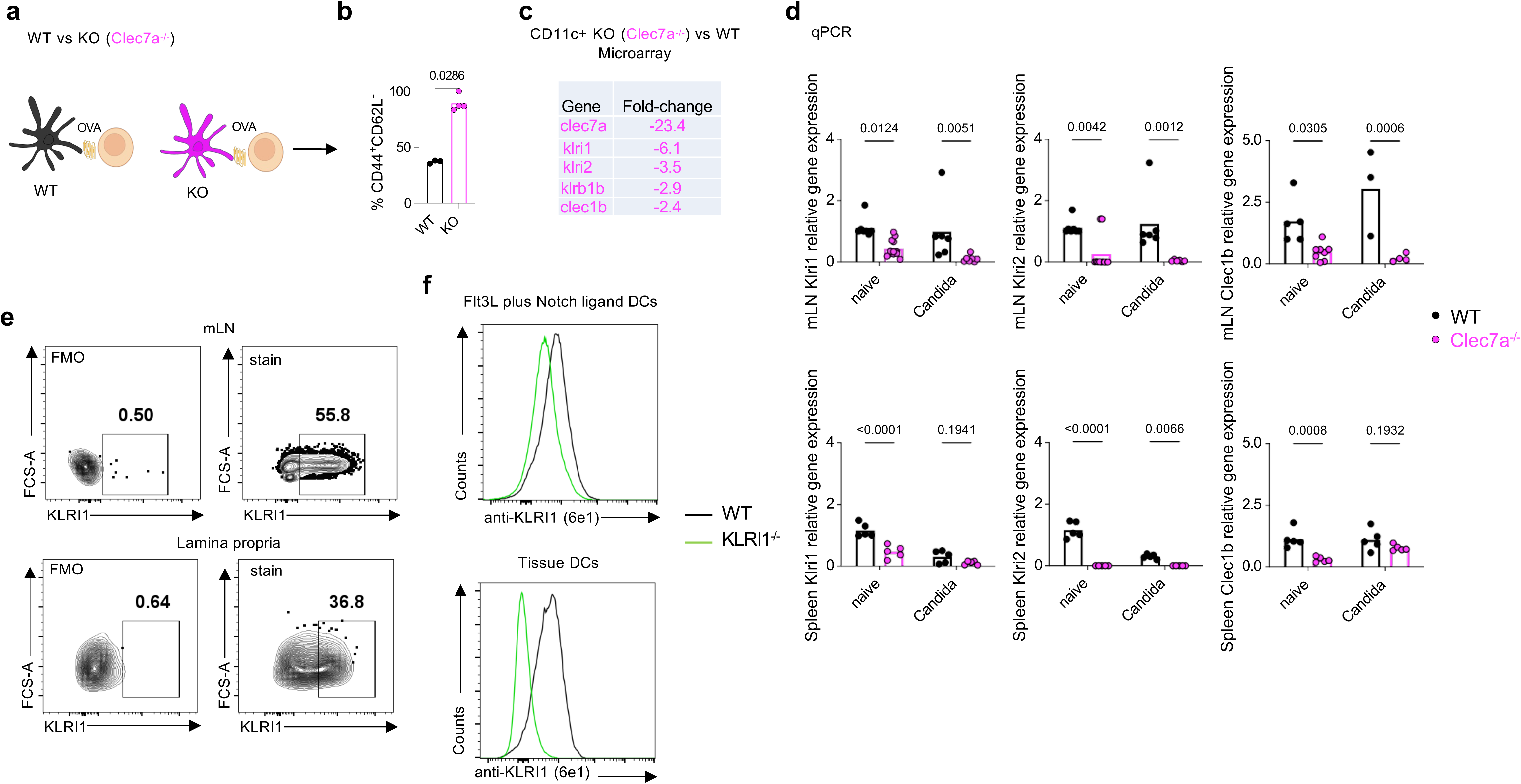
Altered expression of Killer Like Receptors (KLRs) in Clec7a deficient mice. (**a**) DC:T cell co-culture schematic: mLN CD11c⁺ DCs from naïve WT or Clec7a^-/-^ mice co-cultured with naïve (CD44^-^CD62L^+^) OT-II cells in the presence of OVA. (**b**) Frequency of CD44⁺CD62L⁻ OT-II cells analysed 72 hrs post culture. (**c**) Microarray fold-change analysis of gene expression in mLN CD11c^+^ DCs at day 3 post-C. *albicans* infection, showing the most downregulated genes in Clec7a^-/-^ mice compared to WT mice. (**d**) qPCR of *klri1*, *klri2*, and *clec1b* in mLN at steady state and post-C. *albicans* infection. (**e**) PrimeFlow detection of *klri1* transcripts in mLN and gut lamina propria DCs (Lin⁻CD11c⁺MHC-II⁺). (**f**) KLRI1 protein detection with anti-KLRI1 monoclonal antibody (6E1) on FLT3L/Notch-derived BMDCs and tissue DCs. Each point represents one mouse or culture of one experiment, bars indicate mean. Data is representative of three independent experiments. FMO in flow plots indicates fluorescence minus one, and stain refers to full stain. Statistical analysis was performed using Student’s *t*-test for comparing two groups and Two-way ANOVA for more than two groups. p ≤ 0.05 is considered significant.

As we had observed previously^15^, mLN DCs from both naïve and *C. albicans-*infected Clec7a^-/-^ mice displayed relatively minor differences in the expression pattern of B7 and TNFR co-signalling molecules when compared to WT DCs (**Supplementary Fig 2a and b**). This indicates that the significantly enhanced T cell activation observed in Clec7a^-/-^ mice does not primarily stem from the altered expression of classical B7 or TNFR co-signalling molecules.

To gain further insight, we profiled mRNA expression of mLN-derived CD11c^+^ DCs from *C. albicans*-infected Clec7a^-/-^ mice and WT mice. More broadly, we found that loss of Clec7a profoundly reconfigures the transcriptional landscape, with the highest alterations in the expression of other CLRs commonly referred to as killer lectin-like receptors (KLRs)^17^, within the natural killer gene complex (NKC) on mouse chromosome 6^18^ (**Fig. 2c**, **Supplementary Fig. 2c-d**). KLRs comprise a diverse family of cell-surface molecules some of which have been reported to act as key modulators of immune responses^18^. One of the most significantly downregulated genes in Clec7a deficient DCs, *clec1b*, has already been implicated in adaptive immune responses, through regulation of DC migration in the draining lymph nodes^19^. We validated our transcriptomic findings by qPCR analysis of mLN tissues under steady-state conditions or following *C. albicans* infection. Consistent with the transcriptomic data, several KLRs including *klri1*, *klri2*, and *clec1b* were markedly downregulated in the mLN of Clec7a^-/-^ mice, irrespective of infection status (**Fig 2d**). Moreover, qPCR analysis revealed that CD11c^+^ cells were the major cell-type expressing these KLRs in the mLN (**Supplementary Fig. 2e**). The expression of other CLRs within the NKC gene complex neighbouring *clec7a* including *clec9a*, and *clec12b* were unaffected in the Clec7a^-/-^ mice (**Supplementary Fig 2f**). Using PrimeFlow RNA assay, we could directly show *klri1* expression in DCs (Lin^-^CD11c^+^MHC-II^+^) isolated from the mLN and gut lamina propria of WT mice (**Fig 2e**). To exclude the possibility of unintended transcriptional changes in genes neighbouring Clec7a arising from the targeted disruption during generation of our Clec7a knockout strain, we performed qPCR of selected KLR gene using an independently generated Clec7a^-/-^ mouse line^20^. We observed similar patterns of dysregulated KLR gene expression in this independent Clec7a knockout line (**Supplementary Fig. 2g**). Thus, these data indicate that loss of Clec7a results in the selective dysregulation of several receptors in DCs, particularly selected members of the KLR family.

### KLRs are expressed at the DC cell surface

We next explored KLR expression on DCs, focussing on KLRI1 and KLRI2. Using *in vitro* systems, we found that bone-marrow derived DCs (BMDC) generated under GM-CSF and IL-4 conditions^21^ only weakly expressed these receptors, while cultures supplemented with FLT3L^22^ or FLT3L plus Notch ligand^23^, led to significant upregulation of *klri1*, and to a lesser extent *klri2* (**Supplementary Fig. 2h**). Using PrimeFlow we could also detect KLRI1^+^ cells in FLT3L-produced BMDCs (**Supplementary Fig. 2i**). In contrast, repeated attempts to detect KLRI2^+^ cells using this technique were unsuccessful, likely reflecting the low expression levels of this receptor. Furthermore, there were no alterations in expression of *klri1* and *klri2* in FLT3L-derived BMDCs derived from MyD88^-/-^ and TRIF^-/-^ mice, revealing that at basal level, Toll-like receptor signalling is not involved in regulating the expression of the KLRs in DCs (**Supplementary Fig. 2j**).

KLRI1 and KLRI2 have been proposed to require the adaptor protein KLRE1 for their surface expression^24^. We could confirm this observation by generating expression constructs encoding mouse KLRI1 and KLRI2 tagged with an HA epitope and co-expressing them with mouse KLRE1 in NIH3T3 fibroblasts. Surface staining with anti-HA antibody showed that KLRI1 (**Supplementary Fig. 2k**) and KLRI2 (**Supplementary Fig. 2l**) essentially required co-expression with KLRE1 to become available at the cell surface.

As KLRI1 was the most dysregulated KLR in Clec7a^-/-^ mice, we prioritized this receptor for further investigation. To validate surface expression at the protein level, we generated a monoclonal antibody against KLRI1 (**6E1**) (**Supplementary Fig. 2m**). Using this reagent, we demonstrated robust KLRI1 expression on FLT3L and Notch ligand-derived WT BMDCs, as well as on WT tissue Lin⁻CD11c⁺MHC-II⁺ dendritic cells compared to KLRI1 deficient cells or tissue (**Fig. 2f, Supplementary Fig. 2n**). Notably, using a further array of markers, we could show that KLRI1 is expressed at the surface of both cDC1 (Lin^-^CD11c^+^MHC-II^+^Clec9a^+^) and cDC2 (Lin^-^CD11c^+^MHC-II^+^SIRPα^+^) of (**Supplementary Fig. 2o-p**). Thus, these data show that the KLRs are expressed on the surface of DCs.

### KLRI1 modulates effector CD4^+^ T cell activation

To explore whether KLRI1 contributes to the T cell phenotype observed in Clec7a^-/-^mice, we commercially generated a mouse deficient in this receptor (**Supplementary Fig 3a**). We validated *klri1* deletion by flow cytometry and qPCR in mLN and spleen, (see **Fig. 2f**, **Supplementary Fig. 3b**). KLRI1-deficiency did not alter the expression of other key genes including *klri2* and *klre1* (**Supplementary Fig. 3b**). Deletion of KLRI1 did not affect the levels of major immune cell populations in the blood (**Supplementary Fig. 3c-d**), indicating that loss of this receptor had no impact on haematopoiesis.

Next, we sought to determine whether KLRI1 influences CD4^+^ T cell responses, focusing initially on its role in T cell activation. *In vitro*, we found that DCs lacking KLRI1 promoted a significantly higher frequency of CD44^+^CD62L^-^ effector antigen-specific T cells compared to WT DCs (**Fig. 3a-b**), similar to what we had observed in Clec7a^-/-^mice. However, unlike Clec7a-deficient cells, these effector T cells displayed highly elevated levels of CD44 expression (**Fig. 3c**), indicating heightened activation. This data therefore suggests that DC-expressed KLRI1 acts as a negative regulator of T cell activation.

**Fig. 3.**
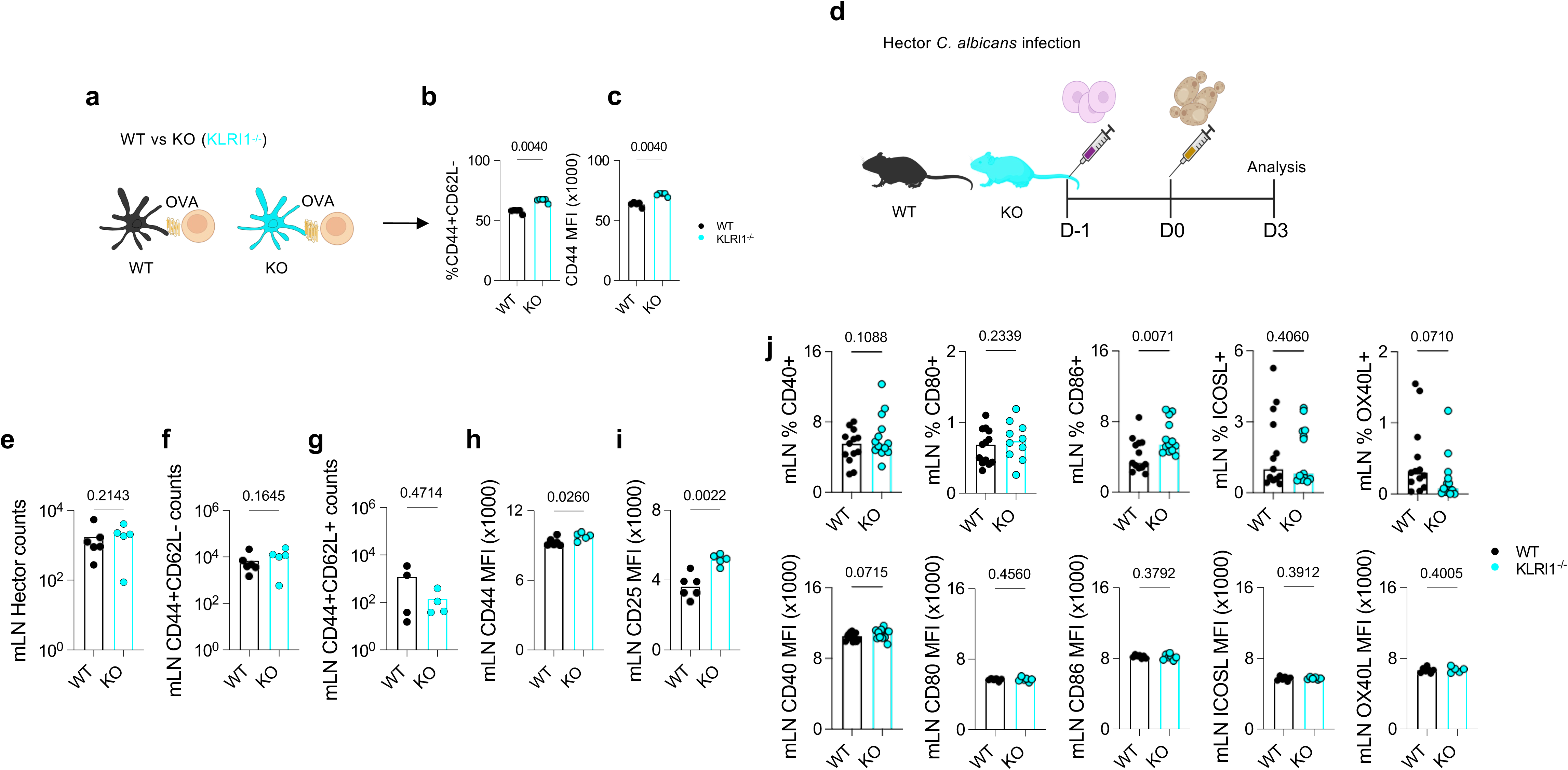
KLRI1 modulates effector CD4^+^ T cell activation. (**a**) DC:T cell co-culture schematic: mLN CD11c⁺ DCs from naïve WT or KLRI1^-/-^ mice co-cultured with naïve (CD44^-^CD62L^+^) OT-II cells in the presence of OVA. (**b**) Frequency of CD44⁺CD62L⁻ OT-II cells analysed 72 hrs post culture and (**c**) CD44 MFI expression of CD44⁺CD62L⁻ OT-II cells. (**d**) Experimental schematic: adoptive transfer of 1×10^6^ naïve Hector CD4^+^ T cells i.v into WT and KLRI1^-/-^ mice at day-1, and next day i.v infection with 1×10^5^ *C. albicans* SC5314 yeasts. mLN flow cytometric analysis of number of live (**e**) *C. albicans*-specific Hector CD4⁺ T cells; (**f**) effector memory CD44^+^CD62L^-^; (**g**) central memory CD44^+^CD62L^+^ with (**h**) CD44 and (**i**) CD25 MFI analysed within the CD44^+^CD62L^-^ subset of Hector cells. (**j**) Flow cytometric analysis of mLN CD11c^+^ DCs from naïve WT and KLRI1^-/-^ mice showing expression of CD40, CD80, CD86, ICOSL, and OX40L. Data depict the frequency of marker-positive cells (top) and their MFI (bottom). Each point represents one mouse; bars indicate the mean. Data are pooled from two independent experiments. Statistical analysis was performed using Student’s *t*-test with p ≤ 0.05 considered significant.

To determine the *in vivo* relevance of KLRI1, we next examined T cell activation in KLRI1^-/-^ mice following systemic *C. albicans* infection (**Fig. 3d**). In the mLN, the number of antigen specific cells was not altered (**Fig. 3e**), nor the number of effector memory CD44^+^CD62L^-^ (**Fig. 3f**) or the CD44^+^CD62L^+^ central memory cells (**Fig. 3g**). However, as we had observed *in vitro*, effector memory CD44^+^CD62L^-^ cells presented with a more activated phenotype in KLRI1^-/-^ mice compared to WT mice, as reflected by their CD44 surface expression levels (**Fig. 3h**). To further substantiate this elevated activation, we assessed expression of CD25, the high-affinity IL-2 receptor α-chain and a canonical marker of T cell activation. Consistent with their heightened activation profile, CD25 level was significantly increased in antigen-specific effector memory CD44^+^CD62L^-^ cells in the absence of KLRI1 (**Fig. 3i**).

We next determined if the enhanced T cell activation in the KLRI1^-/-^ mice could be attributed to changes in classical DC co-stimulatory or co-inhibitory pathways. As we had observed with Clec7a^-/-^, DCs lacking KLRI1 displayed minimal alterations in the expression of co-signalling molecules, both at steady state (**Fig. 3j, Supplementary Fig. 3e**) and following *C. albicans* infection (**Supplementary Fig. 3f**). Together, these findings indicate that KLRI1 is involved in fine-tuning CD4^+^ T cell activation.

### KLRI1 directly binds CD4^+^ T cells and acts as a co-signalling molecule

We then explored the possibility that KLRI1 could directly interact with T cells to modulate their function, and act as a co-signalling receptor. To test this, we first generated Fc-fusion proteins representing homodimeric KLRI1 (FcKLRI1), homodimeric KLRE1 (FcKLRE1), and the heterodimeric KLRE1-KLRI1 complex (FcKLRE1-I1). We then performed a flow cytometry-based binding assay using single-cell suspensions from mLN of naïve mice, in which the CD4⁺ T cell compartment contains both CD44⁺CD62L⁻ effector-memory-like cells and a basal population of CD44⁺CD62L⁺ central memory-like cells^25^. Remarkably, only the heterodimeric FcKLRE1-I1 showed robust and specific binding to CD4^+^ T cells (**Fig. 4a, Supplementary Fig. 4a**). FcKLRI1 and FcKLRE1 homodimers exhibited negligible binding (**Fig. 4a**), indicating that the heterodimer is necessary for CD4^+^ T cell ligand recognition.

**Fig. 4.**
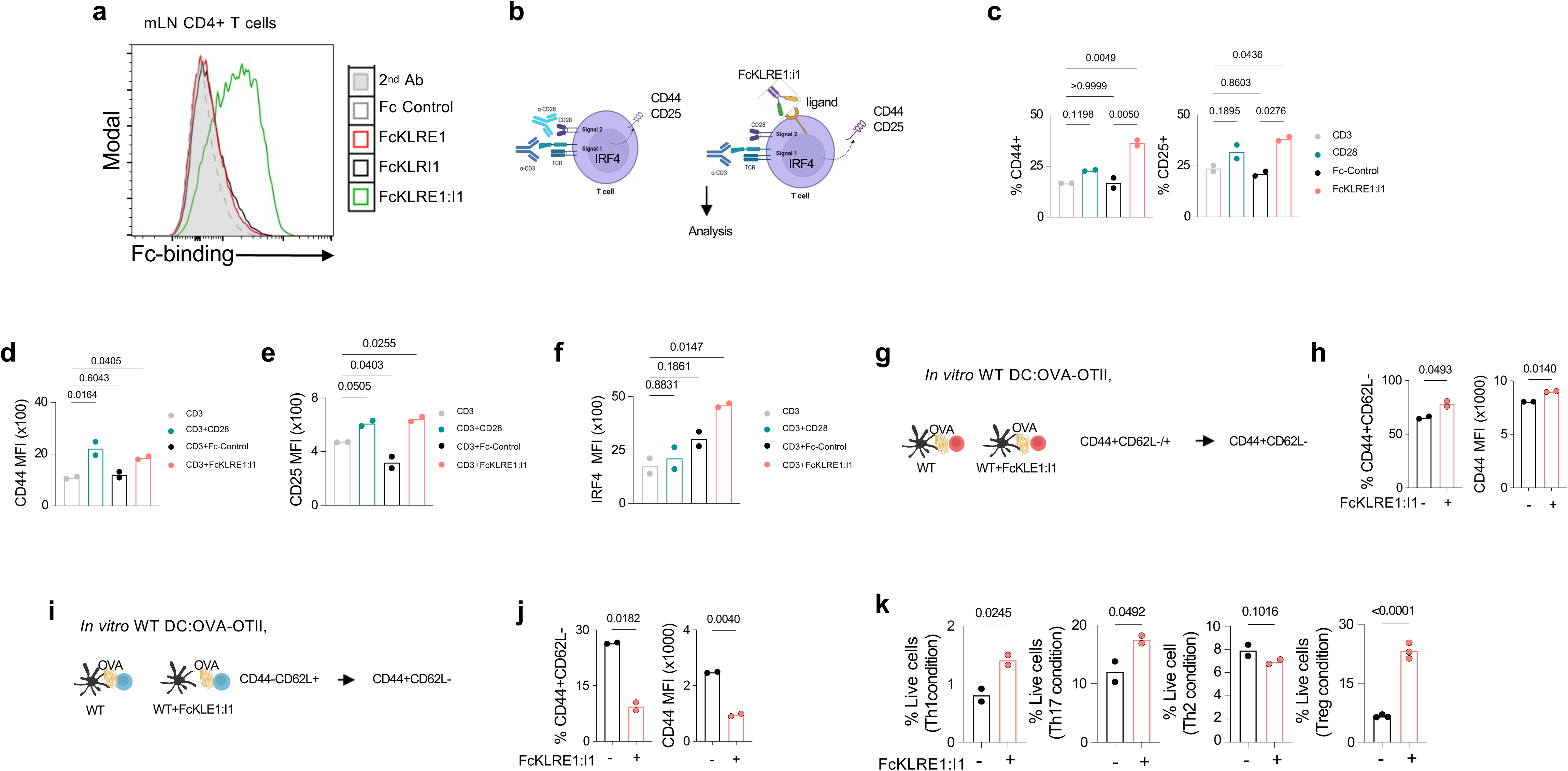
KLRI1 directly binds CD4^+^ T cells and acts as a co-signalling molecule. (**a**) Flow cytometry binding assay of FcKLRI1, FcKLRE1, and FcKLRE1-I1 to CD4⁺ T cells isolated from naïve mLN WT mice. (**b**) Schematic of *in vitro* CD4^+^ T cell activation with plate-bound a-CD3 alone, or with soluble FcKLRE1-I1, or Fc-control, with a-CD28 included as a control. (**c**) Frequencies of activated cells CD44 and CD25, and (**d**) CD44 and (**e**) CD25 MFI levels analysed by flow cytometry 48 hrs following stimulation. (**f**) IRF4 MFI expression after 72 hrs following stimulation with plate-bound a-CD3 alone, or with soluble FcKLRE1-I1, or Fc-control, with a-CD28 included as a control bound. (**g**) DC:T cell co-culture schematic: mLN CD11c⁺ DCs from naïve WT co-cultured with memory (CD44^+^CD62L^-/+^) OT-II cells in the presence of OVA or OVA plus FcKLRE1-I1. (**h**) Flow cytometric analysis of frequency of activated cells (CD44^+^CD62L^-^) and CD44 MFI levels 48 hrs post co-culture. (**i**) DC:T cell co-culture schematic: mLN CD11c⁺ DCs from naïve WT co-cultured with naïve (CD44^-^CD62L^+^) OT-II cells in the presence of OVA or OVA plus FcKLRE1-I1. (**h**) Flow cytometric analysis of frequency of activated cells (CD44^+^CD62L^-^) and CD44 MFI levels 48 hrs post co-culture. (**k**) Frequency of live CD4^+^ T cells under Th1, Th2, Th17, or Treg polarizing conditions with/out FcKLRE1-I1 analysed by flow cytometry 3 days following 2-cycles of polarisation conditions. Each point represents one mouse or culture wells; bars indicate mean. Data are representative of one of three independent experiments. Statistical analysis: Statistical analysis was performed using Student’s *t*-test with p ≤ 0.05 considered significant.

To determine whether the KLRE1-KLRI1 complex can directly modulate CD4^+^ T cell activation as a co-signalling molecule, we assessed its functional impact in a defined *in vitro* system. Specifically, we asked whether KLRE1-KLRI1 engagement could substitute for, or augment, classical co-stimulation in the context of TCR engagement^26^. Using plate-bound α-CD3 antibody to provide TCR cross-linking, we cultured naïve (CD44^-^CD62L^+^) CD4^+^ T cells in the presence of soluble FcKLRE1-I1 heterodimer or an Fc-control protein (**Fig. 4b**). As a positive control, additional wells received soluble α-CD28 to mimic conventional co-stimulation^26^ (**Fig. 4b**). T cell activation was evaluated by quantifying the frequency of activated cells and surface expression of CD44 and CD25 48 hrs following stimulation. As anticipated, stimulation with α-CD3 in combination with α-CD28 increased T cell activation (**Fig. 4c-e**). Strikingly, however, when α-CD3 was provided together with FcKLRE1-I1, CD4^+^ T cell activation was significantly enhanced, as reflected by increased frequency of cells expressing CD44 and CD25 (**Fig. 4c**) as well as stimulating elevated expression of CD44 (**Fig. 4d**) and CD25 (**Fig. 4e**). These results reveal that KLRE1-KLRI1 can function as a potent co-stimulatory signal for T cells, operating independently of the classical CD28 pathway.

To further evaluate the functional consequences of KLRE1-KLRI1 stimulation on CD4^+^ T cells, we assessed expression of IRF4, a transcription factor that acts as a key downstream integrator of TCR and co-stimulatory signals, making it a sensitive marker of productive T cell activation^27^. We show that during TCR cross-linking with α-CD3, FcKLRE1-I1 treatment resulted to a marked increase in IRF4 expression compared to α-CD3 alone or Fc-control (**Fig. 4f**). This data indicates that KLRE1-KLRI1 not only promotes T cell activation but also drives downstream transcriptional programs associated with T cell effector activation.

To gain insight into the expression dynamics of the KLRE1-KLRI1 ligand on CD4^+^ T cells, we analysed FcKLRE1-I1 binding across different T cell maturation states. We found that binding of the FcKLRE1-I1 heterodimer was absent on purified naïve CD44^-^CD62L^+^ CD4^+^ T cells but became detectable approximately 24 hrs after TCR stimulation (**Supplementary Fig. 4b**).

Given that FcKLRE1-I1 binding to CD4^+^ T cells occurs after TCR stimulation, we next investigated whether its functional impact also depends on the maturation state of the responding T cells. To address this, we compared the effects of FcKLRE1-I1 on highly purified naïve (CD44^-^CD62L^+^) versus memory (CD44^+^CD62L^-^ plus CD44^+^CD62L^+^) antigen-specific CD4^+^ T cells from the peripheral lymph nodes of OT-II mice. Both populations were co-cultured with OVA-pulsed DCs derived from WT mLN. This side-by-side comparison revealed a striking dichotomy in how KLRE1-KLRI1 influences CD4^+^ T cell activation. From the purified memory CD4^+^ T cells (**Fig. 4g**), addition of FcKLRE1-I1 significantly enhanced cell activation, as indicated by increased number of effector CD44^+^CD62L^-^ cells as well as of increased CD44 MFI (**Fig. 4h**), as we had observed before. In contrast, when naïve (CD44^-^CD62L^+^) CD4^+^ T cells were stimulated under the same conditions (**Fig. 4i**), FcKLRE1-I1 significantly suppressed their activation relative to control conditions, with decreased number of effector CD44^+^CD62L^-^ cells as well as reduced CD44 MFI (**Fig. 4j**). Thus, these data highlight a context-dependent nature of KLRE1-KLRI1 signalling, functioning as a co-stimulatory axis for antigen-experienced T cells while dampening activation of naïve T cells.

We then explored how KLRE1-KLRI1 influences CD4^+^ T cell responses in the context of TCR cross-linking in the presence of cytokine-driven differentiation. We cultured CD4^+^ T cells under Th1, Th2, Th17, and Treg polarizing conditions in the presence of FcKLRE1-I1. This *in vitro* system allowed us to assess the impact of KLRE1-KLRI1 on T cell survival and impact on CD4^+^ T cell differentiation. FcKLRE1-I1 significantly enhanced CD4^+^ T cell survival under Th1, Th17, and Treg polarizing conditions, but had no significant impact under Th2 conditions (**Fig. 4k**). These findings suggest that KLRE1-KLRI1 selectively promotes the survival and differentiation of Th1-, Th17-, and Treg-lineage CD4^+^ T cells while exerting minimal influence on Th2 polarization. Collectively, these findings position KLRE1-KLRI1 as a context-dependent regulator of T cell activation.

### KLRE1-KLRI1 reconstitution restores anti-fungal immunity and modulates CD4^+^ T cell responses in Clec7a^-/-^ mice

Given that the T cell activation phenotype observed in KLRI1^-/-^ mice resembled that of the Clec7a^-/-^ mice, we hypothesized that the abnormal CD4^+^ T cell responses in Clec7a^-/-^ mice may arise, at-least in part, from a failure to engage KLRI1-mediated co-signalling. To test this, we reconstituted KLRI1 function *in vivo* using the soluble FcKLRE1-I1 in Clec7a^-/-^ mice challenged systemically with *C. albicans* (1×10^5^ yeasts) (**Fig. 5a**). We observed that Clec7a^-/-^ mice treated with FcKLRE1-I1 in the presence of adoptively transferred antigen-specific CD4^+^ T cells exhibited significantly reduced fungal burdens in the GIT, accompanied by restored intestine tissue integrity when compared to untreated control mice (**Fig. 5b, Supplementary Fig. 5a**). This anti-fungal effect was not replicated by treatment with homodimeric FcKLRE1 or FcKLRI1 (**Fig. 5c**), nor by an unrelated Fc-control protein (**Supplementary Fig. 5b**). Furthermore, FcKLRE1-I1 treatment failed to confer protection in the absence of antigen-specific CD4^+^ T cells (**Fig. 5d-e**), confirming that its protective effects are T cell dependent.

**Fig. 5.**
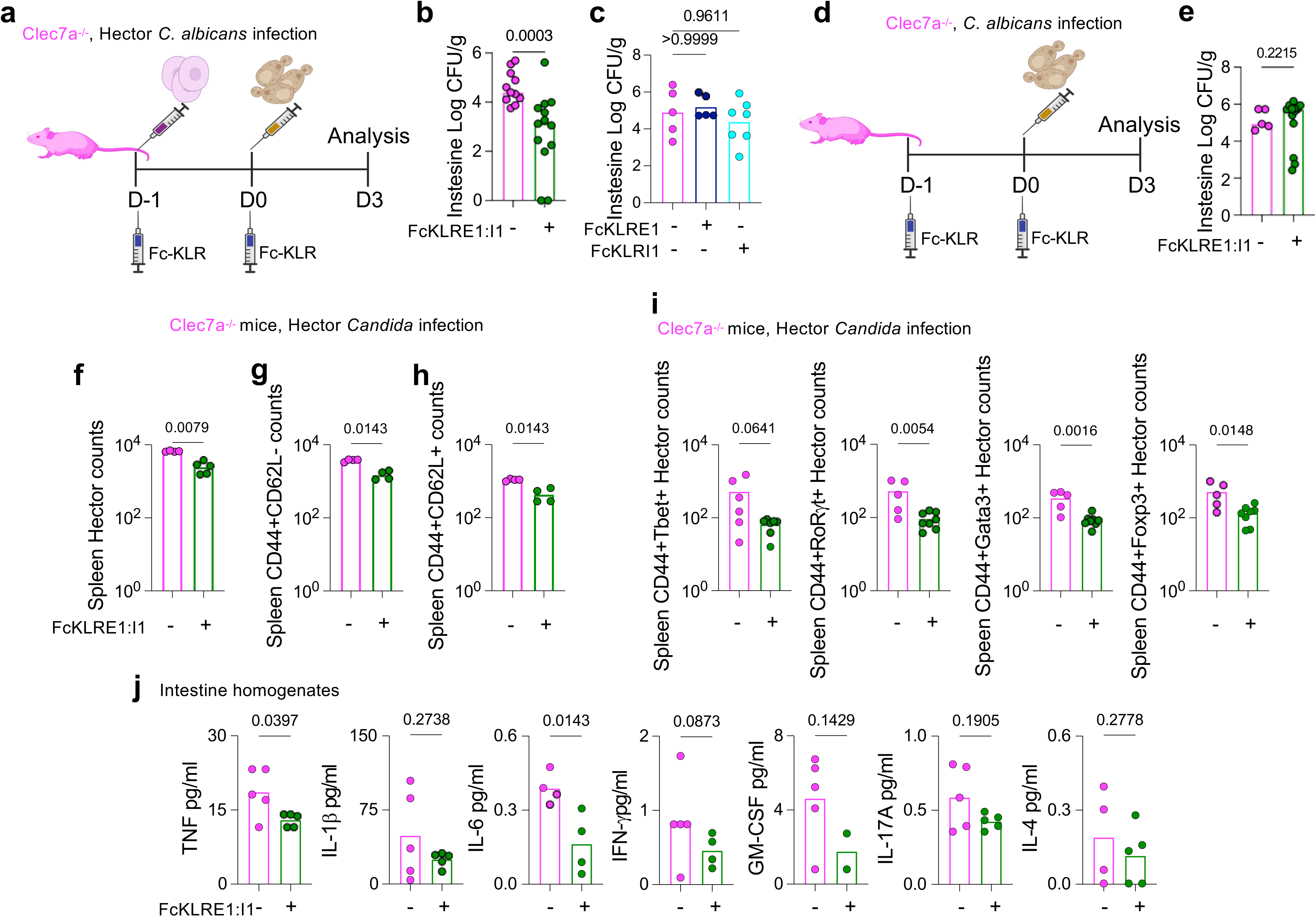
KLRE1-KLRI1 reconstitution restores anti-fungal immunity and modulates CD4^+^ T cell responses in Clec7a^-/-^ mice. (**a**) Treatment and infection schematic of Clec7a^-/-^mice: adoptive transfer of 1×10^6^ naïve Hector CD4^+^ T cells i.v with/out 7.5 ug/ml of FcKLRE1-I1 i.p at day-1, and next day i.v infection with 1-1.5×10^5^ *C. albicans* SC5314 yeasts and a second dose of 7.5 ug/ml of FcKLRE1-I1 i.p. (**b**) Intestine fungal burden 3 days post *C. albicans* infection. (**c**) Fungal burden after treatment with homodimeric FcKLRE1 or FcKLRI1. (**d**) Treatment and infection schematic of Clec7a^-/-^mice with/out 7.5 ug/ml of FcKLRE1-I1 i.p at day-1, and next day i.v infection with 1-1.5×10^5^ *C. albicans* SC5314 yeasts and a second dose of 7.5 ug/ml of FcKLRE1-I1 i.p (**e**) Intestine fungal burden 3 days post *C. albicans* infection. Spleen flow cytometric analysis of number of live (**f**) *C. albicans*-specific Hector CD4⁺ T cells; (**g**) effector memory CD44^+^CD62L^-^; (**h**) central memory CD44^+^CD62L^+^ and (**i**) numbers of antigen-specific CD44^+^Tbet^+^, CD44^+^RORgt^+^, CD44^+^GATA3^+^, and CD44^+^Foxp3^+^ cells. (**j**) TNF, IL-1b, IL-6, IFN-g, GM-CSF, IL-17A and IL-4 concentrations in intestinal tissue homogenates were analysed with Luminex assay. Each point represents one mouse; bars indicate mean. Data shown are representative of three independent experiments, except for Luminex assay that represents a single experiment. Statistical analysis was performed using Student’s *t*-test for two group comparison or Two-way ANOVA for comparisons of three groups with p ≤ 0.05 considered significant.

We next examined the response of CD4^+^ T cells following FcKLRE1-I1 treatment in *C. albicans*-infected mice. In treated Clec7a^-/-^ mice, we noted a significant reduction in overall antigen-specific CD4^+^ T cell counts (**Fig. 5f, Supplementary Fig. 5c**) as well as the effector memory CD44^+^CD62L^-^ (**Fig. 5g, Supplementary Fig. 5d**) and central memory CD44^+^CD62L^+^ antigen-specific cell counts (**Fig. 5h, Supplementary Fig. 5e**). We next examined the polarization of these activated antigen-specific CD4⁺ T cells, focusing on the master transcription factors Tbet, RORγt, Gata3 and Foxp3, which define Th1, Th17, Th2 and Tregs lineages, respectively. This analysis was prompted by previous reports showing elevated Th1-and Th17-associated cytokine responses in Clec7a^-/-^ mice compared with WT mice following *C. albicans* infection^12, 13^. Treatment of Clec7a^-/-^ mice with FcKLRE1-I1 resulted in a significant reduction of antigen-specific CD44^+^Tbet^+^, CD44^+^RORγt^+^, CD44^+^RORγt^+^, CD44^+^Gata3^+^, and CD44^+^Foxp3^+^ cell counts (**Fig. 5i, Supplementary Fig. 5f**), compared to untreated Clec7a^-/-^ mice. We also assessed cytokine profiles in intestinal tissue homogenates by Luminex assay. Relative to untreated Clec7a^-/-^ mice, TNF, IL-1β, IL-6, IFN-γ, GM-CSF, IL-17A, IL-4 displayed a downward trend with TNF and IL-6 reaching significance following FcKLRE1-I1 treatment in Clec7a^-/-^ mice (**Fig. 5j**). These data indicate that the addition of FcKLRE1-I1 to Clec7a^-/-^ mice restores tissue integrity, modulate effector CD4^+^ T cell and cytokine balance leading to improved control of systemic *C. albicans* infection.

## Discussion

Clec7a is an archetypal C-type lectin receptor (CLR) known for its essential role in anti-fungal immunity through β-glucan recognition. It signals via the Syk-CARD9 axis to drive phagocytosis, inflammasome activation, and pro-inflammatory cytokine production^4^. However, an expanding body of literature reveals that Clec7a functions far beyond fungal resistance, contributing to immunity against bacteria, viruses, and parasites, and even impacting diseases including colitis, asthma, neuroinflammation, and tumor immunity^28, 29^. These pleiotropic effects are especially evident in CD4^+^ T cell-dependent responses, where Clec7a has been linked to shaping Th1 and Th17 immunity, but the molecular underpinnings of the global influence beyond its innate immune sensing functions remain elusive. Our study sheds light on the underlying mechanisms, helping to interpret these broad phenotypes.

The aberrant CD4^+^ T cell phenotype in Clec7a^-/-^ mice, marked by hyperactivation, apoptosis, and failure to control fungal growth, has been previously attributed to altered innate cytokine output or impaired antigen presentation^2, 15, 37, 38^. We identify a transcriptional axis downstream of Clec7a that governs the expression of a large subset of KLRs within the NKC locus in dendritic cells. We explored the most strongly Clec7a regulated gene, *klri1*, whose role in CD4 immunity is poorly understood. We discovered that KLRI1, in heterodimeric complex with KLRE1, is a context-dependent co-signalling receptor that shapes CD4^+^ T cell activation and survival. We show that the ligand for KLRE1-KLRI1 is not constitutively expressed on CD4^+^ T cells but emerges after TCR engagement, mirroring the activation-dependent behaviour of other co-stimulatory receptors such as ICOS^30^, OX40^31^, and the inhibitory checkpoint PD-1^32^. Notably, we found that KLRE1-KLRI1 acts as a context-dependent modulator of T cell activation: enhancing responses in memory T cells while suppressing activation of naïve CD4^+^ T cells. This dichotomy is reminiscent of PD-1 and CTLA-4, which dampen priming of naïve cells but allow more nuanced regulation in effector and memory compartments^33^. Functionally, KLRE1-KLRI1 engagement upregulates IRF4, a transcriptional integrator of TCR signal strength that guides effector lineage decisions and metabolic fitness^34, 35^. Remarkably, reintroduction of KLRE1-KLRI1 signaling into Clec7a^-/-^ mice through FcKLRE1-I1 administration restored balanced CD4^+^ T cell activation and improved anti-fungal control during systemic *C. albicans* infection. This effect was strictly dependent on the presence of antigen-specific CD4^+^ T cells with FcKLRE1-I1 treatment also limiting broad effector CD4^+^ T cell differentiation, reducing the number of cells that express T-bet (Th1), RORγt (Th17), GATA3 (Th2), and Foxp3 (Treg). In parallel, intestinal cytokine levels associated with both innate (TNF, IL-1β, IL-6) and adaptive (IFN-γ, GM-CSF, IL-17A, IL-4) immune responses were decreased in FcKLRE1-I1 treated Clec7a^-/-^ mice. Together, these findings support the notion that FcKLRE1-I1 recalibrates immune activation in Clec7a^-/-^ mice, dampening excessive inflammatory signaling while maintaining effective anti-fungal resistance and preserving tissue integrity. This immune response dampening resembles function of checkpoint molecules such as LAG-3, TIGIT, and BTLA, which safeguard tissues from immune pathology in high-antigen or chronic inflammatory settings^36^.

The OVA-CpG and OVA-curdlan settings, performed in the absence of live pathogens, reveal that Clec7a deficiency mediates a broader defect in CD4^+^ T cell immunity. In the presence of cognate antigen, Clec7a^-/-^ mice exhibited markedly increased numbers of activated antigen-specific CD4⁺ T cells, which we show is in part due to absence of KLRI1 expression in Clec7a^-/-^ mice. These findings demonstrate that the KLRE1-KLRI1 complex operates as a co-signaling checkpoint modulating CD4^+^ T cell activation not only during *C. albicans* infection but also under sterile inflammatory conditions. The mirrored phenotypes across infectious and non-infectious models underscore the robustness of this regulatory axis and reveal that Clec7a’s influence on T cell activation is not pathogen-restricted but rooted in its transcriptional control of key co-signaling molecules such as KLRI1.

The KLRE1-KLRI1 axis exerts its regulatory influence, at least in part, through an inducible ligand expressed on CD4^+^ T cells. The ligand’s expression pattern suggests that its availability is tightly coupled to the T cell activation state. Indeed, ligand detection only after TCR engagement implies that activated T cells themselves may provide a transient, self-limiting checkpoint cue analogous to PD-L1/PD-1 or Tim-3/Galectin-9 interactions^33, 39, 40^. This configuration would enable CD4^+^ T cells to modulate their own effector output and maintain homeostasis within inflamed tissue. This observation positions the KLRE1-KLRI1:KLRE1-KLRI1 ligand pair as a dynamic, activation-sensitive checkpoint within the adaptive compartment, which is poised to dampen excessive T cell activation once antigen-specific responses are underway. Whether the KLRE1-KLRI1 ligand is unique to CD4^+^ T cells or expressed by other leukocyte subsets remains an open avenue for discovery.

In summary, our data establish Clec7a not only as an innate β-glucan sensor but also as a transcriptional gatekeeper that controls expression of non-canonical KLRs with checkpoint-like properties. The Clec7a-KLR axis that we have identified likely underlies previously unexplained CD4⁺ T cell phenotypes observed in Clec7a^-/-^ models of infection, autoimmunity, asthma, neuroinflammation, and gut homeostasis. In the future, defining the CD4^+^ T cell ligand for KLRE1-KLRI1 and delineating its downstream signaling pathways will be essential to fully elucidate how this checkpoint mediates its functions in T cell activation or inhibition and preserves immune homeostasis. Furthermore, testing these pathways in DC-specific Clec7a conditional knockouts will be essential. In parallel, we will extend our analysis to additional KLRs uncovered in our datasets, including molecules like KLRI2, to define how this broader network shapes adaptive responses (see associated manuscript, Salazar et al^43^). Finally, determining how KLRI1 interfaces with classical checkpoints such as PD-1 or CTLA-4 may open new avenues for combinatorial immunotherapy. By uncovering an innate-adaptive checkpoint axis governed by Clec7a, we redefine this CLR as a central co-ordinator of immune homeostasis with implications extending well beyond anti-fungal resistance.

## Methods

### Mice

C57BL/6J were purchased from Charles River Laboratories, UK. Clec7a^-/-^ were previously generated^2^. KLRI1^-/-^ mice (full knockout mouse on C57BL/6N background) was generated using CRISPR/Cas9-mediated genome engineering (Cyagen, USA) (**Supplementary Fig 3a**). Briefly, mouse genomic fragments containing homology arms were amplified from a BAC clone using high-fidelity Taq DNA polymerase and sequentially assembled into a targeting vector together with recombination sites and selection markers. Exon 1 was selected as the conditional knockout region. Guide RNA (gRNA) targeting the *klri1* gene and Cas9 mRNA were co-injected into fertilized mouse zygotes to generate knockout offspring. F0 founder mice were identified by PCR and confirmed by sequence analysis. The line was subsequently rederived onto a C57BL/6N background at Charles River Laboratories, UK and were bred and maintained under specific pathogen-free (SPF) conditions. Hector transgenic^16^ and OT-II transgenic mice were maintained under SPF conditions at the University of Exeter, UK. Mice were co-housed for at least 14 days before experiments and used at 6-8 weeks of age. For all studies, age-matched females were randomly assigned to experimental or control groups. Femurs and tibia from MyD88^-/-^ and TRIF^-/-^ mice were kindly provided by Andrew S. MacDonald (University of Edinburgh, UK). Spleens from Clec7a^-/-^ mice generated using a different approach^20^ were kindly provided by Shinobu Saijo (Chiba University, Japan). All procedures complied with the University of Exeter ethical review process and UK Home Office licence PP9965358.

### Adoptive transfer of OT-II or Hector CD4⁺ T cells

Single-cell suspensions from peripheral lymph nodes (pLNs) of OT-II (CD45.1) or Hector (CD90.1) mice were prepared in PBS. CD4^+^ T cells were purified by negative selection using the EasySep Mouse CD4^+^ T Cell Isolation Kit (STEMCELL Technologies), as per manufacturer’s instructions, washed twice, counted using a Vi-Cell (Beckman Coulter), and adjusted to 10×10⁶ cells/ml. A total of 1×10⁶ cells in 100 µl PBS was injected intravenously (i.v.) (lateral tail vein) into mice.

### Systemic *Candida albicans* infection

Mice were challenged intravenously with either 1×10⁵ or 2-2.5 x 10⁵ yeasts of *C. albicans*. For Hector cell studies, mice were infected with the wild-type *C. albicans* strain SC5314 24 hours (hrs) after adoptive transfer. For OT-II cell studies, mice were infected 24 hrs after adoptive transfer with the OVA-expressing *C. albicans* strain Calb-Ag^41^. Yeasts were grown in YPD broth at 30 °C for 24 hrs, washed twice in PBS, counted, and adjusted to the required concentration. For KLR reconstitution studies, Clec7a^-/-^ received two intraperitoneal injections of purified Fc–fusion protein (10 µg in 100 µl PBS): the first 24 hrs before *C. albicans* infection and the second immediately prior to infection. At the indicated time points, mice were euthanised and tissues collected. For fungal burden, organs were homogenised in PBS, serially diluted, and plated on YPD agar supplemented with gentamicin (100 µg/ml) and vancomycin (10 µg/ml). Plates were incubated at 37 °C for 24 hrs before counting CFU.

### Non-infection models

OT-II recipients were injected intraperitoneally with 100 ul of 100 μg ovalbumin (OVA; InvivoGen) plus either 50 μg CpG oligodeoxynucleotides (class X; InvivoGen) in PBS or 2 mg curdlan (β-1,3-glucan; InvivoGen) freshly sonicated in PBS.

### Preparation of single-cell suspensions

Mesenteric lymph nodes (mLNs), pLNs and spleens were processed through 70 µm filters into RPMI-1640 GlutaMAX. Peripheral blood was collected into EDTA-coated tubes. Lamina propria cell were isolated using the lamina propria dissociation kit (Miltenyi Biotec) according to manufacturer’s instructions. Red blood cells were lysed using BD PharmLyse (BD Biosciences), followed by washing (400 x g, 5 min) and resuspension in RPMI-1640 GlutaMAX with 10 % FCS and 1 % penicillin/streptomycin (complete RPMI) for downstream assays.

### RNA extraction, cDNA synthesis and quantitative PCR

RNA from purified cells or tissues was extracted with TRIzol (Thermo Fisher Scientific) as per manufacturer’s instructions, quantified on a NanoDrop (Thermo Fisher Scientific), and reverse-transcribed using SuperScript III (Thermo Fisher Scientific) as per manufacturer’s instructions. qPCR was performed on a QuantStudio 7 Pro system (Thermo Fisher Scientific), using PowerTrack SYBR Green Master Mix (Thermo Fisher Scientific) as per manufacturer’s instructions. Cycling: 95 °C for 2 minutes (min); 40 cycles of 95 °C for 5 seconds (s), Tm 60 °C for 30 s, followed by melt curve analysis. Relative expression was calculated by the ΔΔCt method normalised to Gapdh and 18srrna. The following primers were used: Klre1 (forward 5′-GTCAGTTTGTTCGCACAG-3′; reverse 5′-TGAAACTCGCAGACAGTG-3′), Klri1 (forward 5′-GGTAACTGGCATGCTTGGAG-3′; reverse 5′ CAAAGATCACAGGAGGCGTC-3′), Klri2 (forward 5′-ACCCTCGAAGACAGACCAGAG-3′; reverse 5′-CCACATCGAATCCTGTCCTG-3′), Klrb1b (forward 5′-GCATGATTCACCTCCATCTC-3′; reverse 5′-GATAGCACCAGACACAGAG-3′), Clec1b (forward 5′-GTTGCTACGGGTTCTTCAG-3’; reverse 5′-CCTTTCTGCAATGTAGTCCAG-3’), Klrc1 (forward 5′-TGAAGGTGGCAAAGAACTC-3’; reverse 5′-AGTCCGAATAGATGATTTCCTG-3’), Clec7a (forward 5′-TTAGACTTCAGCACTCAAGAC-3’; reverse 5′-ACTACTACCACAAAGCACAG-3’), Clec1a (forward 5′-ACCATGCTGAAGATAAGCAC-3’; reverse 5′-GAGCCCTGTCCAATAAGAG-3’), Klrd1 (forward 5′-CTTCTCCAACCACCACTG-3’; reverse 5′-ATTTCTGGATTGGGGCTG-3’), Olr1 (forward 5′-CAATCTTTGGGTGGCCAG-3’; reverse 5′-CCGATGCAATCCAATCCAG-3’), Gapdh (forward 5′-ACGACCCCTTCATTGACCTC-3′; reverse 5′-CATTCTCGGCCTTGACTGTG-3′) and 18srrna (forward 5′-ATGAACGAGGAATTCCCAG-3′; reverse 5′-CCAATCGGTAGTAGCGAC-3′).

### Flow cytometry

Single-cell suspensions were prepared in PBS containing 2 % FCS and 2 mM EDTA (FACS buffer), stained with viability dye eFluor 780 (eBioscience), washed, then stained with fluorochrome-conjugated monoclonal antibodies for same-day acquisition or fixed with 2 % PFA (Sigma). Conjugated antibodies included: anti-CD80-FITC/BV650 (clone 16-10A1, BioLegend), anti-CD40-BV605 (clone 3/23, BioLegend), anti-CD86-AF700 (clone GL1, eBioscience), anti-CD275-PE (ICOSL; clone HK5.3, supplier BioLegend), anti-CD252-APC (OX40L; clone RM134L, BioLegend), anti-CD90.1-BB700/R718 (clone HIS51, BD Biosciences), anti-CD90.1-APC (clone OX-7, BD Biosciences), anti-CD45.1-BV480 (A20, BD Biosciences), anti-CD45-BUV496/PerCP-Cy5.5 (clone 30-F11, BD Biosciences/BioLegend), anti-CD4-BUV563/BV510/BUV496 (clone RM4-5/RM4-4, BD Biosciences/BioLegend), anti-CD3-AF594/eFluor 450 (clone 500A2, BioLegend/eBioscience), anti-CD44-FITC/BV510/PE-Cy5 (clone IM7, BD Biosciences/BioLegend), anti-CD62L-BV605/PE/BUV563 (clone MEL-14, BD Biosciences), anti-B220-eFluor 450/AF700 (clone RA3-6B2, eBioscience/BioLegend), anti-CD49b-eFluor 450 (clone DX5, eBioscience), anti-CD11b-BUV395 (clone M1/70, BD Biosciences), anti-CD335-SB645 (NKp46; clone 29A1.4, eBioscience), anti-CD11c-BV711 (clone HL3, BD Biosciences), anti-I-A/I-E-BUV496 (MHC-II; clone 2G9, BD Biosciences), anti-XCR1-APC/BV510 (clone ZET, BioLegend), anti-CD172a-PerCP-eFluor710 (SIRPα; clone P84, eBioscience), anti-CD26-BUV737 (clone H194-112, BD Biosciences), anti-CD64-AF647 (clone X54-5/7.1, BD Biosciences), anti-Ly6G-eFluor450/SB550 (clone 1A8, eBioscience/BioLegend), anti-Ly6C-BV570 (clone HK1.4, BioLegend), anti-CD170-SB436 (Siglec-F; clone 1RNM44N, eBioscience), anti-CD25-BV650/PE-Cy7/PE/FITC/PerCP-Cy5.5 (clone PC61, BD Biosciences/BioLegend), anti-F4/80-FITC (clone BM8, BioLegend), anti-CX3CR1-BV785/PE (clone SA011F11, BioLegend), anti-CD161-AF647/BV605 (NK1.1; clone HP-3G10/PK136, BioLegend/BD Biosciences), anti-IRF4-PE/BV786 (clone Q9-343, BD Biosciences), anti-T-bet-BV421/BV786 (clone O4-46, BD Biosciences), anti-GATA3-PE-Cy7/PE-CF594 (clone L50-853, BD Biosciences), anti-FoxP3-AF488 (clone MF23, BD Biosciences), and anti-RORγt-BV650 (clone Q31-378, BD Biosciences). Intracellular staining was conducted using the Transcription Factor Staining Buffer Set (eBioscience) as per manufacturer’s instructions. Annexin V (BioLegend) staining was performed as per manufacturer’s instructions. For Fc–protein staining, cells were incubated with Fc–fusion proteins (100 μg/ml) for 60 min at 4 °C, washed, and stained with anti-human IgG-PE (Jackson ImmunoResearch). For anti-KLRI1 staining, cells were blocked with 5% mouse serum and incubated with biotinylated primary antibodies, washed, and labelled with streptavidin conjugated anti-mouse IgG-APC (Jackson ImmunoResearch). Data were acquired on a Attune NxT (Thermo Fisher Scientific) or Cytek Aurora and analysed using FlowJo v10 (BD Biosciences).

### Generation and purification of Fc–fusion proteins

The ectodomain of Klre1 and Klri1 were amplified from murine spleen cDNA with primers Klre1 (forward 5′-GTCAGTTTGTTCGCACAG-3′; reverse 5′-TGAAACTCGCAGACAGTG-3′) and Klri1 (forward 5′-GGTAACTGGCATGCTTGGAG-3′; reverse 5′-CAAAGATCACAGGAGGCGTC-3′) containing BamHI and NotI sites using Q5 High-Fidelity DNA polymerase (New England Biolabs) as per manufacturer’s instructions. Amplicons were gel-purified (Wizard SV Gel and PCR Clean-Up System, Promega), cloned into Zero Blunt TOPO vectors (Invitrogen) as per manufacturer’s instructions, sequenced, and subcloned into pSecTag2a-Fcmut. HA (KLRE1) or 6×His (KLRI1) tags were added by overlap-extension PCR.

### Transfection

HEK293T cells were maintained in complete DMEM medium (Gibco; high glucose, GlutaMAX™, pyruvate), supplemented with 10% FBS (Gibco; Fetal Bovine Serum, heat inactivated); 20mM HEPES buffer solution (Gibco) and 100U/ml Penicillin-Streptomycin (Gibco), at 37°C with 5% CO2. One day before transfection, HEK293T cells with >95% viability, were prepared at 106 viable cells/ml in 75cm2 flasks. Plasmid DNA was linearised with FastDigest NsbI restriction enzyme (Thermo Scientific) and purified by column (Qiagen; QIAquick PCR Purification Kit). Stable transfection was achieved with FuGENE® HD reagent (Promega) at 1:6 ratio (transfection reagent:DNA), following manufacturer instructions, with 5µg in total linearised plasmid DNA added per transfection. For the expression of the heterodimers, DNA from each construct was added at 1:1 ratio to the cells. After transfection, cells were incubated at 37°C for 2 days before starting selection with 400µg/ml Zeocin (InvivoGen) added to the complete medium. Transfected HEK293T cells were lifted with TrypLE Express (Gibco) and transferred to 175cm2 flask with selective medium, incubated at 37°C for two hours and then at +4°C for another two hours, before continuing growth in selective media for several weeks, until resistant clones were established.

### Protein expression

Once the transfected cell line was established, protein expression was achieved by growing the cells in 875 cm2 cell surface multi-flasks (Falcon™), in DMEM medium supplemented with 5% FBS, without zeocin selection. After 4 days growth, supernatant from the cells was collected, cleared of cell debris by centrifugation at 2000xg and filtered through a 0.22µm filtering unit (Nalgene). The presence of the DNA sequence for KLR protein expression in the transfected cells was confirmed by PCR amplification from genomic DNA extracted from the zeocin resistant clones, following manufacturer instructions for the genomic DNA prep with dilution buffer (Thermo Fisher; Phire Tissue Direct PCR Kit). PCR amplification with DreamTaq DNA Polymerase (Thermo Fisher) with initial denaturation at 95°C for 2min, followed by 30 cycles denaturation 30sec at 95°C, annealing 30sec at 51°C and elongation 1min at 72°C, with a final extension at 72°C for 5 minutes. Genomic DNA prep diluted 1:100 in water before amplification with primers T7 (5’ - TAATACGACTCACTA TAGGG – 3’) and FcR2 (5’ - GTTGAACTTGACCT CAGGGTC – 3’). Protein expression was confirmed by ELISA assay to detect the presence of the Fc chimeric proteins. Maxisorp ELISA plates (Thermo Fisher) were coated with 100µl of supernatant from transfected HEK293T cells and incubated over-night (O/N) at +4°C. Plates were washed with 1x PBS + Tween20 (0.05%) [PBS-T] and blocked with 200µl of 10% FBS in 1x PBS for 2 hours. Plates were washed with PBS-T and 100µl peroxidase-conjugated anti-Human IgG (Jackson ImmunoResearch), at 1/10000 dilution, was added to the wells. After 1 h incubation at room temperature, wells were washed with PBS-T and 100µl of TMS substrate (ThermoFisher) was added to develop the reaction. Reaction was stopped with 50µl 2N H2SO4 and optical densities measured at 450nm.

### Protein purification

Soluble protein in the supernatant of transfected HEK293T cells was purified using Protein A sepharose (Cytiva; rProtein A Sepharose Fast Flow resin) by gravity flow. For the heterodimers (KLRE1/KLRi1; KLRE1/KLRi2), Ni Sepharose 6 Fast Flow (Cytiva; HisTrap FF) and Anti-HA Agarose (Pierce™) was used to purify protein fraction with both proteins present. Anti-HA agarose was used for protein purification by gravity flow using a disposable 5ml column (Thermo Scientific). HisTrap FF 1ml column was used according to manufacturer recommendations (wash buffer with 25mM Imidazole; elution with 500mM Imidazole). Ab Buffer Kit (Cytiva) was used for Protein A and HA-column purification, following manufacturer recommendations. Purified protein was dialyzed into 1xPBS (Gibco) using dialysis cassettes (Thermo Scientific™ Slide-A-Lyzer™ G3 Dialysis Cassettes, 10K MWCO). Empty columns and purification accessories were treated with 1M NaOH for two hours, then rinsed with PBS, before protein purification.

### Bone marrow-derived dendritic cells (BMDCs)

Femurs and tibia were aseptically removed, cleaned, and flushed with RPMI using a 25G needle. Suspensions were passed through 70 µm strainers and red blood cells lysed with BD PharmLyse (BD Biosciences). For GM-CSF/IL-4 cultures, bone marrow cells were plated at 1×10⁶/ml in DMEM-GlutaMAX with 10 % FCS, 1 % penicillin/streptomycin, 50 µM 2-mercaptoethanol, GM-CSF (20 ng/ml; BioLegend) with or without IL-4 (10 ng/ml; Bio-Techne), with fresh medium added on day 3. For Flt3L-derived DCs, 4×10⁶/ml bone marrow cells were cultured in complete RPMI-1640 containing 10 % FCS, 1 % penicillin/streptomycin, 50 µM 2-mercaptoethanol and Flt3L (200 ng/ml; BioLegend), with fresh medium added on day 3. For OP9-supported DCs, Flt3L-day 3 cells (2×10⁶/ml) were seeded on confluent mitomycin-C treated (Stell Cell Technologies) OP9 monolayers expressing the Notch ligand Delta-like 1 (DLL1) using complete RPMI in the presence of Flt3L. BMDCs were collected at day 7-8 for further analysis. OP9 cells were maintained in α-MEM with 20 % FCS, 1 % penicillin/streptomycin.

### PrimeFlow RNA assay

Single-cell suspensions (2×10⁶ cells) were stained for surface markers, fixed, permeabilised, and hybridised with Klri1 (Alexa Fluor 647), probe sets (Thermo Fisher Scientific) as per manufacturer’s instructions. Data were acquired on a Attune NxT (Thermo Fisher Scientific) and analysed in FlowJo v10.9.

### T cell activation and co-culture assays

Purified CD4^+^ T cells were stimulated with plate-bound α-CD3 (1 ug/ml) and soluble α-CD28 (2 μg/ml; 2B Scientific), with or without FcKLR proteins (10 μg/ml) for 24-72 hrs. For DC-T cell co-cultures, mLN-derived DCs isolated using CD11c^+^ MicroBeads UltraPure (Miltenyi Biotec) as per manufacturer’s instructions were pre-incubated with OVA or alcohol dehydrogenase (ADH) peptides (2 μg/ml) before adding OT-II or Hector T cells (5:1 DC:T cell ratio) with or without FcKLR proteins (10 μg/ml). Activation markers were analysed by flow cytometry as above.

### T cell polarisation

CD4^+^ T cells were purified from pLN as described above and resuspended in complete RPMI. Cells (200 μl per well) were stimulated with plate-bound α-CD3 (1 ug/ml) and soluble α-CD28 (2 μg/ml; 2B Scientific), in the presence of cytokines and blocking antibodies to direct lineage commitment. For Th1 differentiation: IL-12 (10 ng/ml; Bio-Techne) with anti-IL-4 (2.5 μg/ml; Bio-Techne). For Th2 differentiation: IL-4 (10 ng/ml; Bio-Techne) with anti-IFN-γ (2.5 μg/ml; Bio-Techne). For Th17 polarisation: IL-6 (50 ng/ml; Bio-Techne), TGF-β (2 ng/ml; Bio-Techne), IL-1β (10 ng/ml; Bio-Techne), anti-IFN-γ (2.5 μg/ml; Bio-Techne) and anti-IL-4 (2.5 μg/ml; Bio-Techne). For T regulatory cell induction: TGF-β (10 ng/ml; Bio-Techne). For multiple polarisation cycles, cells were maintained for 96 hrs with daily supplementation of IL-2 (10 ng/ml; Bio-Techne) before re-stimulation under the same polarising conditions.

### Microarray analysis

RNA samples were sequenced using the Ion Torrent Proton sequencer (Centre for Genome Enabled Biology and Medicine, University of Aberdeen Genomics facility). Raw fastq files were successively processed in the following order through Fastqc (v. 10.1), Trimgalore (v. 3.1), Samtools (v. 1.19), STAR aligner (v. 2.4) and Htseq (v. 5.4). Genome alignment was conducted against the Mus musculus_ NC_000072.7 chromosomes file provided by the Gene expression analysis was performed using Partek® Genomics Suite® software, version 6.6 Copyright ©: 2015. Absolute cut-off value was set at 1.5-fold change in expression (KO vs. WT). The statistical significance limit was set at 0.05. K-mean clustering, with applied restriction to 3 clusters and a false discovery rate set at 0.05 was performed via the STRING resource (V. 12) and the Cytoscape v. 3 Clue GO plugin (Bindea et al., 2009). Network construction was performed with Cytoscape V.3 freeware. Statistical comparison among GO term enrichment percentages, and fold-expression bar charts was performed with GraphPad Prism (v. 10.2).

### Generation of NIH-3T3 cells overexpressing KLRs

The coding region of Klre1, Klri2 and Klri1 were amplified from murine spleen cDNA with primers Klre1 (forward 5′-TTTGGATCCACCACCATGGATGAAGCACCTG-3′; reverse 5′-AGCGGCCGCTACTTCTTGCAAATATATGTCAG-3′), Klri2 (forward 5′-GGATCCACCATGCACAAGAAAAAGCATATTAAAC-3′; reverse 5′-CCCCTCGAGTATATTAAATTCACAGATGTATG-3′) and Klri1 (forward 5′-GGATCCACCATGCTTCACAGTAAACGCCGGG-3′; reverse 5′-CCCCTCGAGTATATTAAATTCACAGATGTATG-3′) containing BamHI and XhoI sites for Klri1 and Klri2, and BamH1 and Not1 for Klre1 using Q5 High-Fidelity DNA polymerase (New England Biolabs) as per manufacturer’s instructions. Amplicons were gel-purified (Wizard SV Gel and PCR Clean-Up System, Promega), cloned into Zero Blunt TOPO vectors (Invitrogen) as per manufacturer’s instructions, sequenced, and subcloned into pFb-Neo-HA vector for Klri1 and Klri2 and pMXs-IP vector for Klre1 (subcloned into pFb-Neo-HA vector and pMXs-IP vector are detailed in our previous paper)^42^. Constructs were purified (Wizard SV Gel and PCR Clean-Up System, Promega) and transfected into Plat-E cells with FuGENE-6 (Promega) as per manufacturer’s instructions. Viral particles were harvested 48 h post-transfection and stored at −80 °C in the presence of polybrene (5 µg/ml). NIH3T3 cells were transduced, and stable clones were selected using Geneticin (400 µg/ml) for pFb-Neo-HA transfectants, or puromycin (1 µg/ml) for pMXs-IP transfectants. After 2 weeks of selection, stable expression of KLR constructs was confirmed by flow cytometry using an anti-HA tag antibody (16B12, Enzo life scientific) and anti-KLRE1-AF488 (854929, Bio-Techne).

### Hybridoma generation and monoclonal antibody production

To generate anti-KLRI1 monoclonal antibody (6e1), KLRI1^-/-^ mice were immunised subcutaneously with 100 µg FcKLRI1 emulsified in TiterMax, followed by three boosters of the same mixture. Serum titres were monitored by ELISA. The highest responder received an intraperitoneal boost with NIH-KLRI1-HA cells before splenocytes were harvested and fused with NS1 myeloma cells (ATCC) using PEG1500 (Sigma). Hybridomas were selected in HAT medium (Sigma), screened by flow cytometry, and high-affinity IgG clones were expanded. Antibody-producing cells underwent three rounds of single-cell cloning by limiting dilution, after which antibodies were purified by affinity chromatography with Protein G resin (Pierce), eluted (glycine-HCl, pH 2.5), neutralised (1 M Tris-HCl pH 9), and dialysed into PBS.

### Luminex multiplex cytokine and chemokine analysis

Cytokines (TNF, IL-1β, IL-6, IFN-γ, IL-17A, GM-CSF, IL-4) were quantified using Bio-Plex Pro Mouse Cytokine kits (Bio-Rad) as per manufacturer’s instructions. Plates were read on a MAGPIX system (Luminex), and concentrations were calculated in a Bio-Plex Manager software.

### Statistical analysis

Statistical tests were performed using GraphPad Prism 10. Data were analyzed using student *t*-tests or Two-ANOVA where more than groups were analysed with p ≤ 0.05 considered statistically significant.

## Acknowledgements

We thank the staff of the animal facilities at the University of Exeter for the care and support of our animals. We acknowledge funding from the MRC Centre for Medical Mycology at the University of Exeter (MR/N006364/2 and MR/V033417/1), the NIHR Exeter Biomedical Research Centre, and the Wellcome Trust (217163/Z/19/Z). Additional work may have been undertaken by the University of Exeter Biological Services Unit. The views expressed are those of the author(s) and not necessarily those of the NIHR or the Department of Health and Social Care. For the purpose of open access, the author has applied a CC BY public copyright licence to any Author Accepted Manuscript version arising from this submission.

---

**Supplementary Fig. 1.**
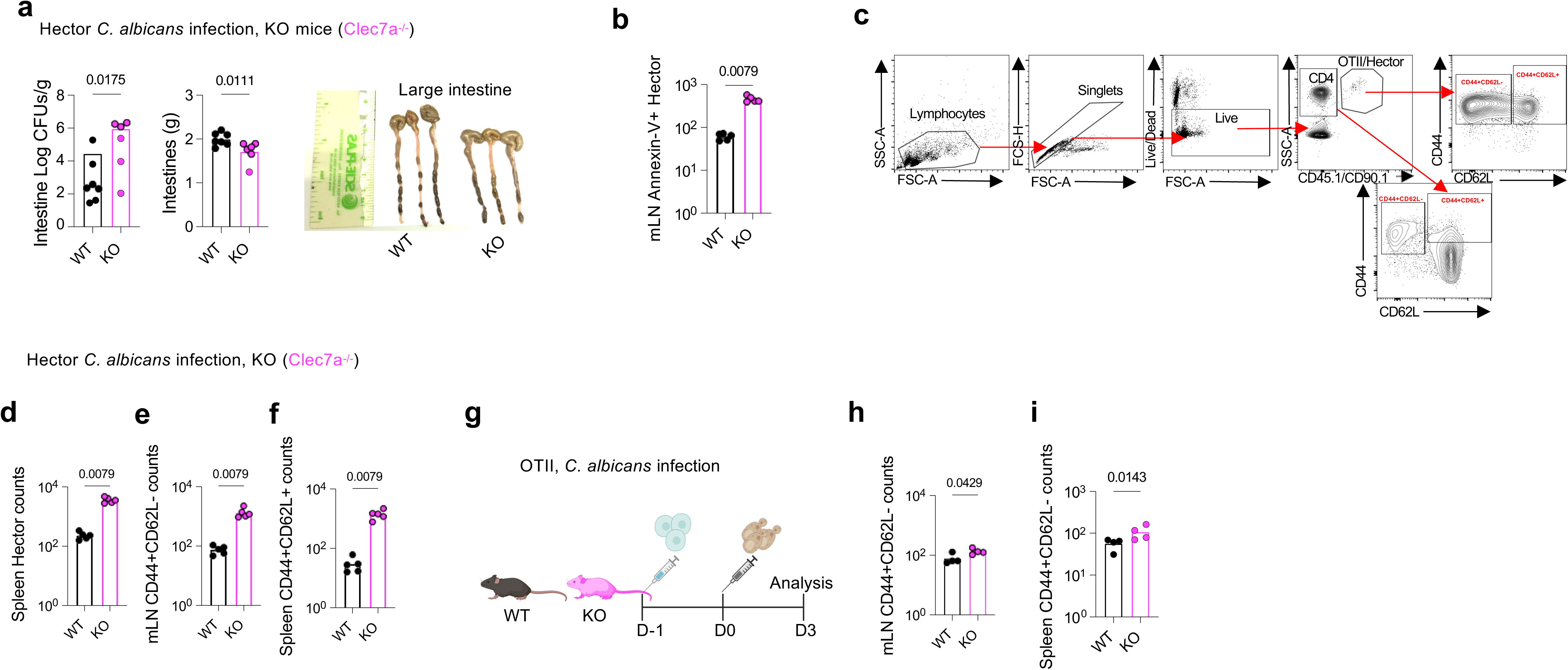
(**a**) Intestinal fungal burden and tissue mass following adoptive transfer of 1×10^6^ naïve Hector CD4^+^ T cells i.v into WT mice and Clec7a^-/-^mice at day-1, and next day i.v infection with higher inoculum of 2-2.5×10^5^ *C. albicans* SC5314 yeasts. Large intestines from WT mice and Clec7a^-/-^mice we imaged at day 3 post infection. (**b**) mLN flow cytometric analysis of apoptotic Hector CD4⁺ T cells. (**c**) Flow cytometry gating strategy to identify live CD4 cells bearing CD90.1 (Hector cells in *C. albicans* model) or CD45.1 (OT.II cells in OVA-CpG or OVA-curdlan models). The effector memory cells were identified as CD44^+^CD62L^-^ and central memory cells are CD44^+^CD62L^+^ cells. Spleen flow cytometric analysis of number of live (**d***) C. albicans-*specific Hector CD4⁺ T cells; (**e**) effector memory CD44^+^CD62L- and (**f**) central memory CD44^+^CD62L^+^ cells. Experimental schematic: adoptive transfer of 1×10^6^ naïve OT-II CD4^+^ T cells i.v into WT mice and Clec7a^-/-^ mice at day-1, and next day i.v. infection with 1×10^5^ OVA-expressing *C. albicans* (Calb-Ag) yeasts. Flow cytometric analysis of number of live effector memory CD44^+^CD62L-from (**h**) mLN and (**i**) and spleen. Each point represents one mouse, and horizontal bars indicate the mean. Data are representative of one of three independent experiments. Statistical analysis was performed using Student’s *t*-test with p ≤ 0.05 considered significant.

**Supplementary Fig. 2.**
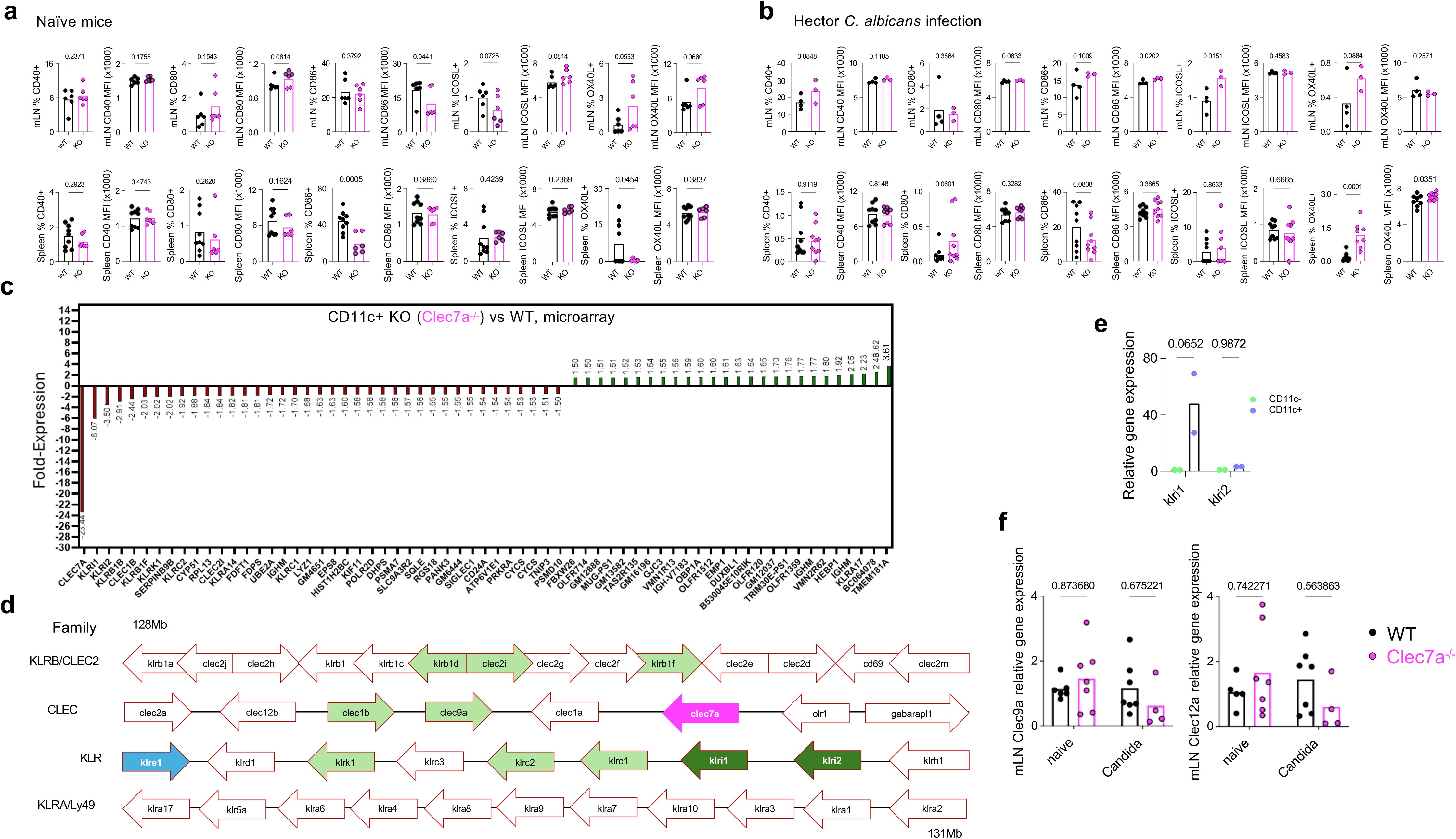
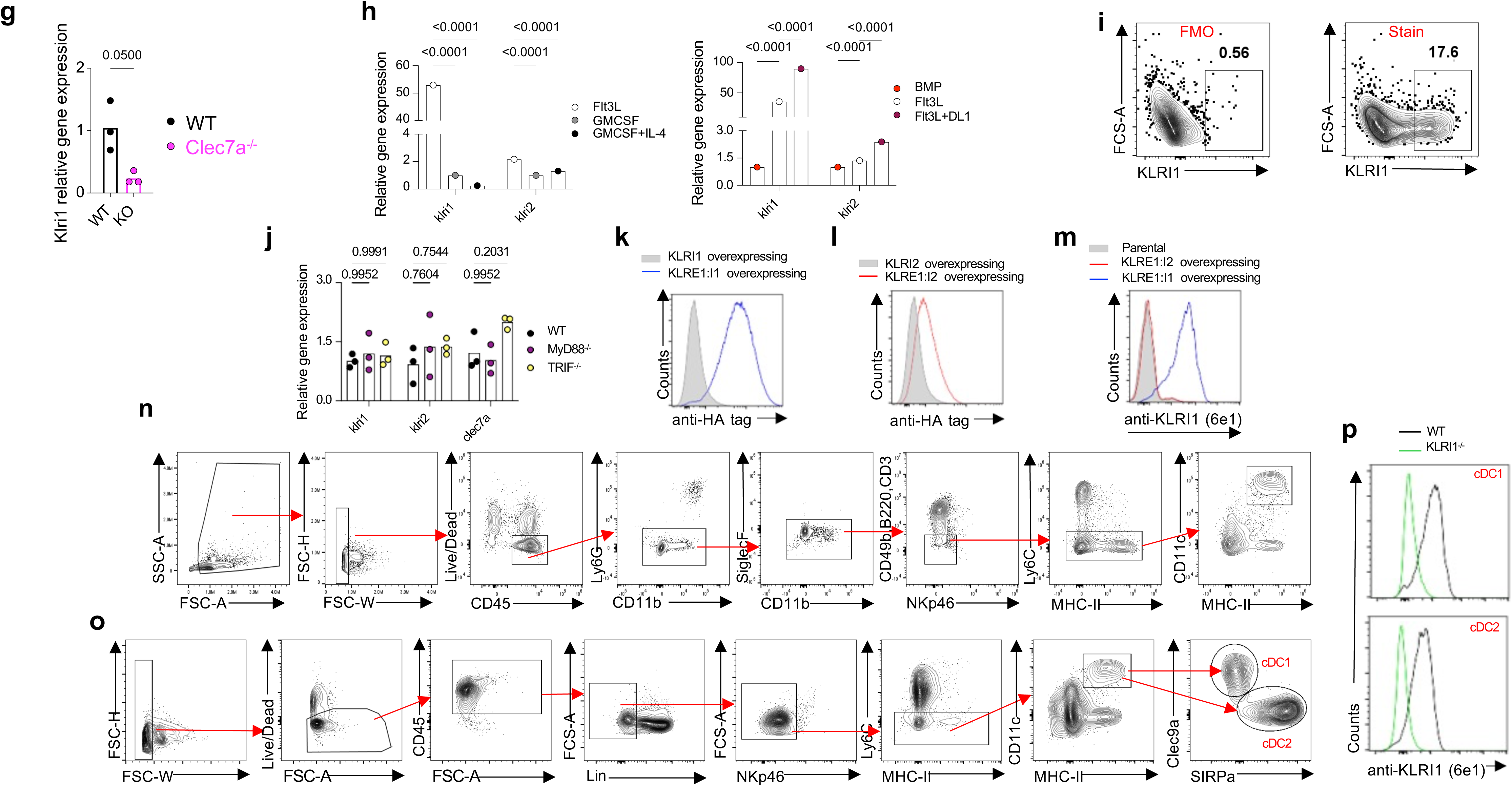
Flow cytometric analysis of frequency of DCs expressing CD40, CD80, CD86, ICOSL, OX40L in DCs as well as their MFI levels in mLN and spleen of (**a**) naïve WT and Clec7a^-/-^ mice and (**b**) WT and Clec7a^-/-^ mice 3 days post *C. albicans* infection. (**c**) Microarray fold-change analysis of gene expression in mLN CD11c^+^ DCs at day 3 post-C. *albicans* infection, showing an extended list the most downregulated or upregulated genes in Clec7a^-/-^ mice compared to WT mice. (**d**) Gene orientation of KLRs within the natural killer gene complex (NKC) on mouse chromosome 6. (**e**) KLRI1 and KLRI2 gene expression in CD11c^+^ and CD11c-cells isolated from the mLN of naïve WT mice. Data are expressed relative to CD11c-cells and normalised to housekeeping genes. (**f**) qPCR of *clec9a* and *Clec12b* transcript levels in mLN of WT and Clec7a^-/-^ mice at steady state and 3 days post *C. albicans* infection. Data are expressed relative to naive controls and normalised to housekeeping genes. Each point represents one mouse; horizontal bars indicate the mean of one experiment. Data are representative of two independently performed experiments. qPCR gene expression of (**g**) *kIri1* analysed from spleen of second Clec7a^-/-^ mouse line; (**h**) *kIri1* and *kIri2* analysed from bone marrow-derived cells cultured for at least 7 days under conditioning media with various cytokines. Data are shown as expression relative to GM-CSF (left) or bone marrow cells (BMP) (right) samples and normalized to housekeeping genes. Bars represent the mean of one experiment performed in triplicates. (**i**) PrimeFlow analysis of KLRI1 expression in FLT3L plus Notch ligand. Left panel shows fluorescent minus one (FMO) control and right panel shows full stain. Data shown is representative of three independent experiments. (**j**) qPCR analysis of *klri1*, *klri2* and *clec7a* gene expression in Flt3L-derived BMDCs from WT, MyD88^-/-^ and TRIF^-/-^. Data are expressed relative to WT mice and normalised to housekeeping genes. Each point represents one mouse; horizontal bars indicate the mean of pooled data from one experiment. (**k**) Flow cytometry analysis of KLRI1 expression in NIH3T3 cell lines overexpressing different constructs. KLRI1 expression was assessed with a specific anti-HA tag monoclonal antibody. Data shown is representative of three independent experiments. (**l**) Flow cytometry analysis of KLRI2 expression in NIH3T3 cell lines overexpressing different constructs. KLRI2 expression was assessed with a specific anti-HA tag monoclonal antibody. Data shown is representative of three independent experiments. (**m**) Flow cytometry analysis of KLRI1 expression in NIH3T3 cell lines overexpressing different constructs. KLRI1 expression was assessed with a specific anti-KLRI1 monoclonal antibody (6e1). Data shown is representative of three independent experiments. (**n**) Flow cytometry gating strategy CD11c⁺MHC-II⁺ for analysis of anti-KLRI1 monoclonal antibody (6e1) on DCs. Flow cytometry analysis gating strategy (**o**) of KLRI1 expression in (**p**) spleen cDC1 (Lin^-^CD11c^+^MHC-II^+^Clec9a^+^) or cDC2 (Lin^-^CD11c^+^MHC-II^+^SIRPα^+^). Data shown are representative of two independent experiments. Statistical analysis was performed using Student’s *t*-test for two group comparison or Two-way ANOVA for comparisons of three groups with p ≤ 0.05 considered significant.

**Supplementary Fig. 3.**
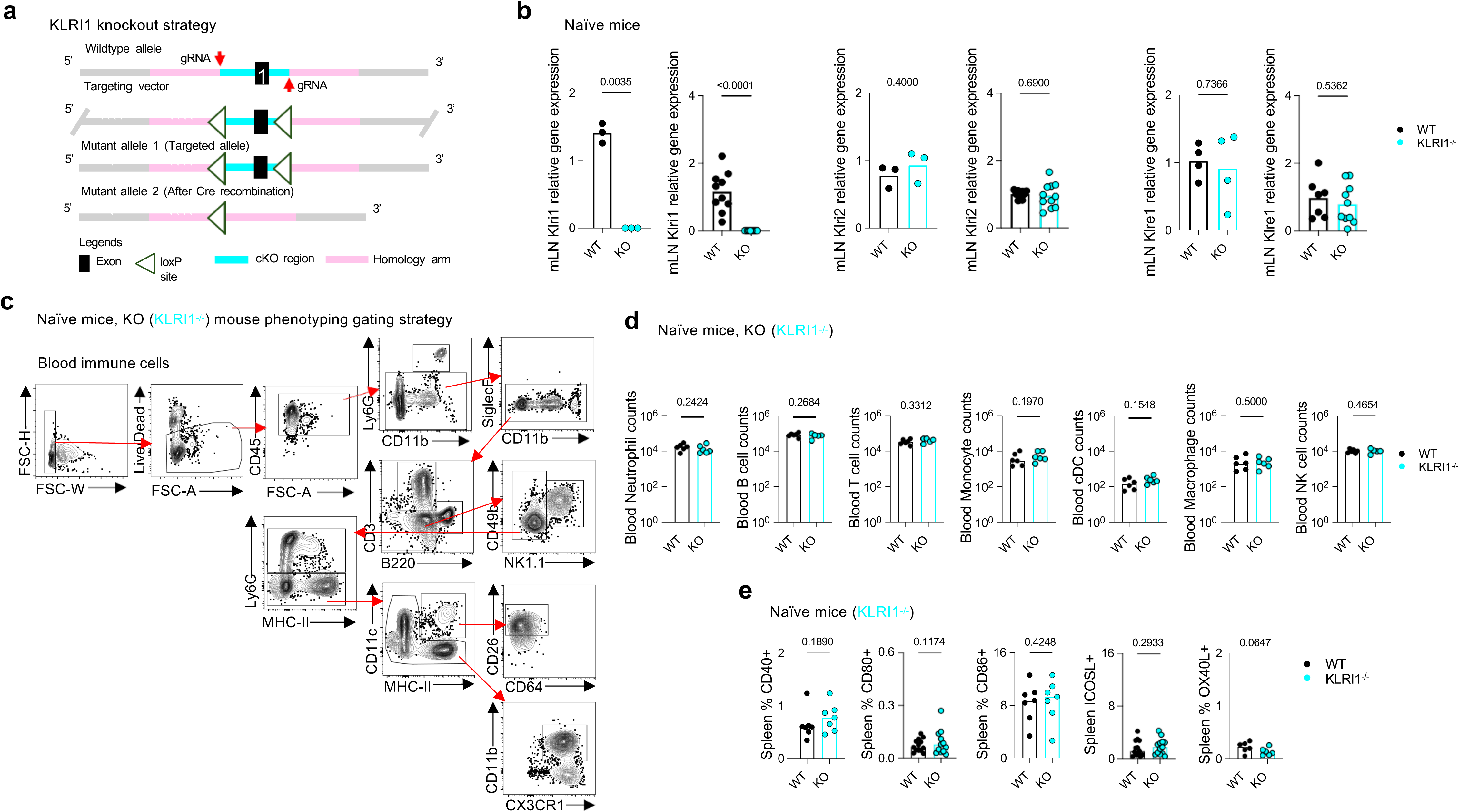
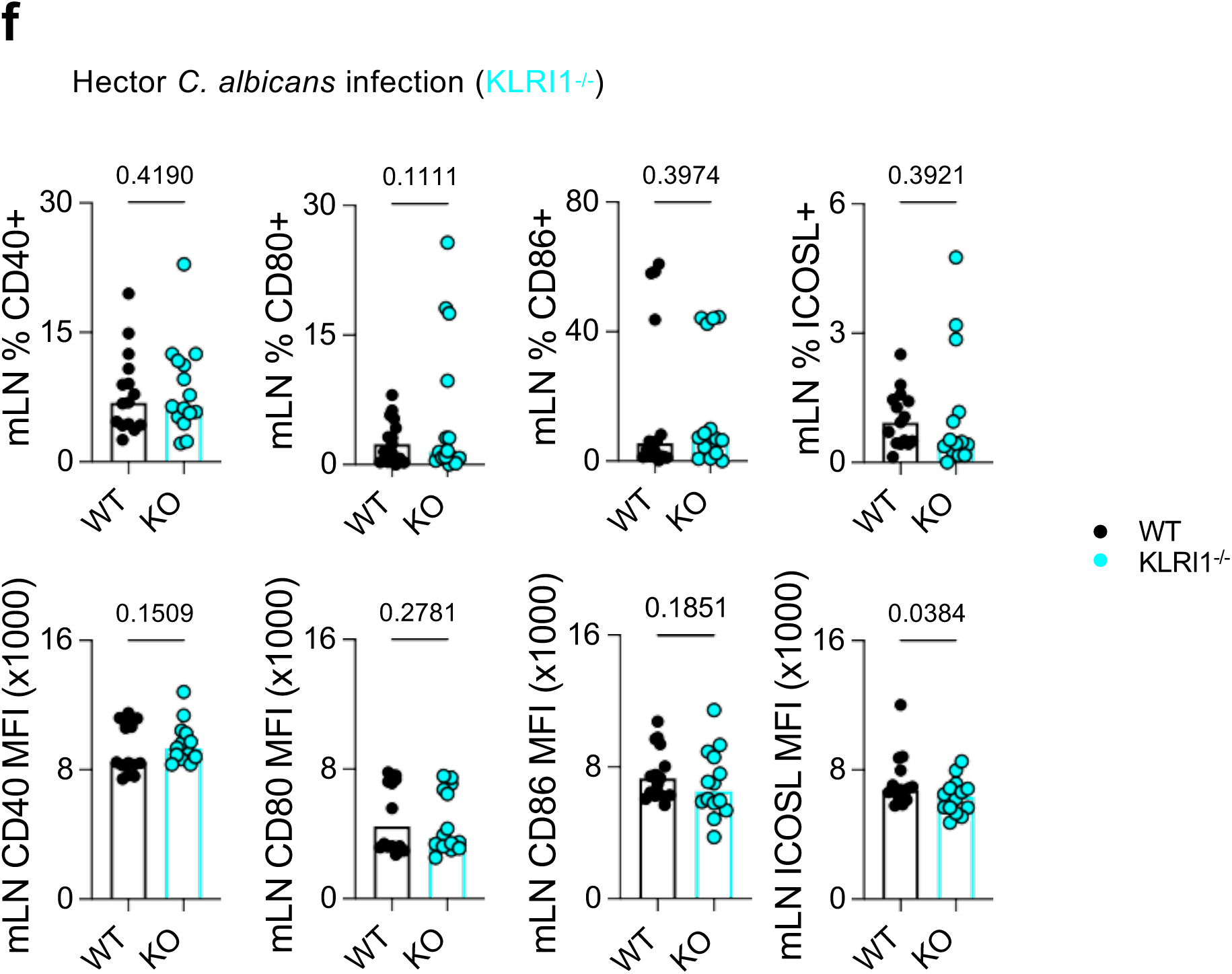
(**a**) Strategy to generate KLRI1-deficient mice. (**b**) qPCR validation of *klri1* deletion in mLN and spleen, and fold-change analysis of *klri2* and *klre1* transcripts from naïve WT and KLRI1^-/-^mice. Flow cytometry (**a**) gating strategy of (**d**) blood immunophenotyping and of naïve WT and KLRI1^-/-^ mice. Flow cytometry analysis of (**e**) splenic CD11c^+^MHC-II^+^ DCs from WT and KLRI1^-/-^ mice expressing co-signalling molecule at steady state. Each point represents one mouse; bars indicate mean. Data are pooled from two independent experiments. Statistical analysis was performed using Student’s *t*-test with p ≤ 0.05 considered significant. (**f**) Flow cytometry analysis of splenic CD11c^+^MHC-II^+^ DCs from WT and KLRI1^-/-^ mice expressing co-signalling molecule from mLN 3 days post *C. albicans* infection. Each point represents one mouse; bars indicate mean. Data are pooled from two independent experiments. Statistical analysis was performed using Student’s *t*-test with p ≤ 0.05 considered significant.

**Supplementary Fig 4.**
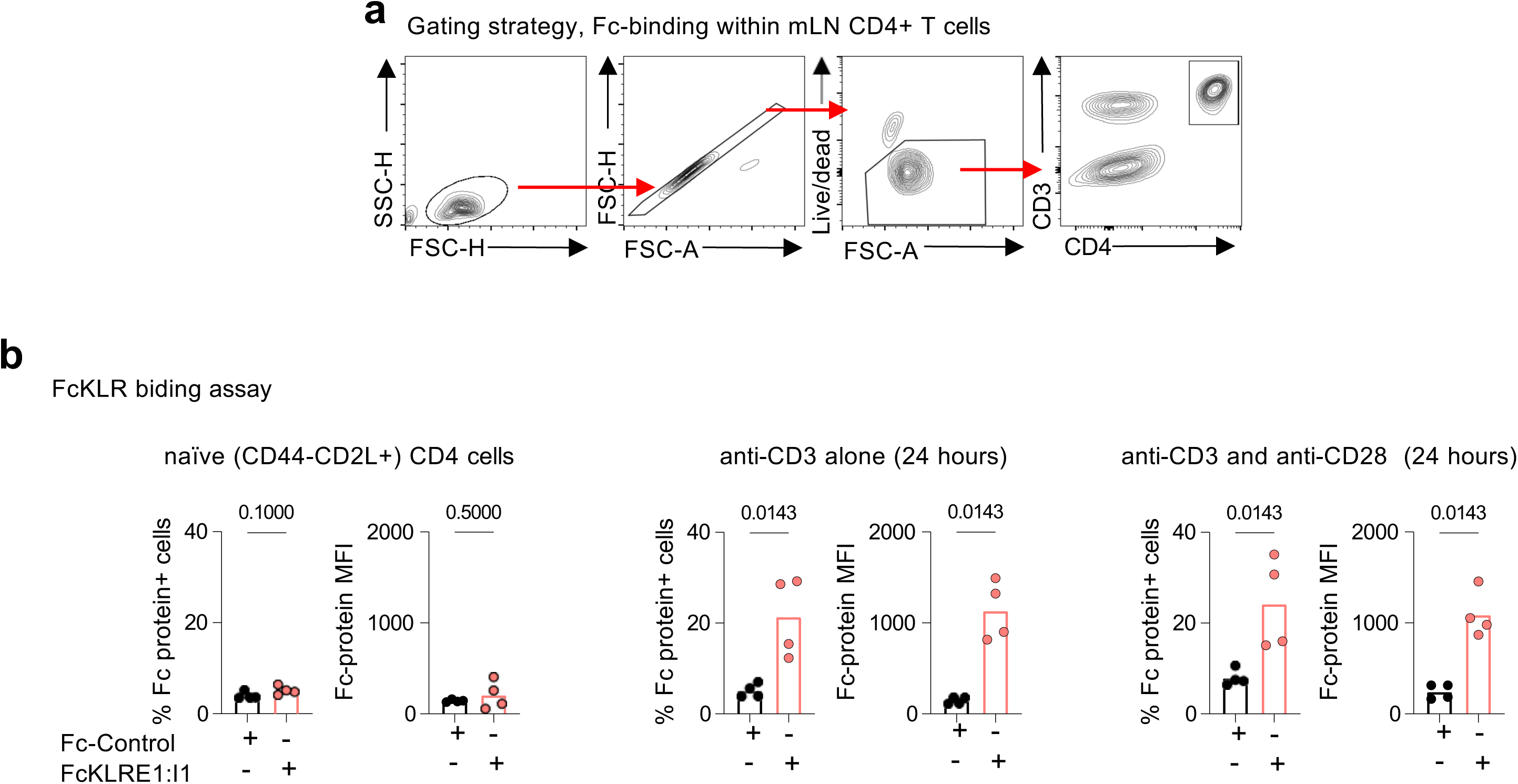
(**a**) Flow cytometry gating strategy for analysing Fc-binding to CD4^+^ T cells. (**b**) FcKLRE1-I1 binding on naïve (CD44^-^CD62L^+^) CD4^+^ T cells or after 24 hrs TCR stimulation with a-CD3 alone, or a-CD3 plus a-CD28 assessed by flow cytometry; with Fc-control used as unrelated KLR molecule. Each point represents one mouse; bars indicate mean. Data are representative of one of four independent experiments. Statistical analysis: Statistical analysis was performed using Student’s *t*-test with p ≤ 0.05 considered significant.

**Supplementary Fig. 5.**
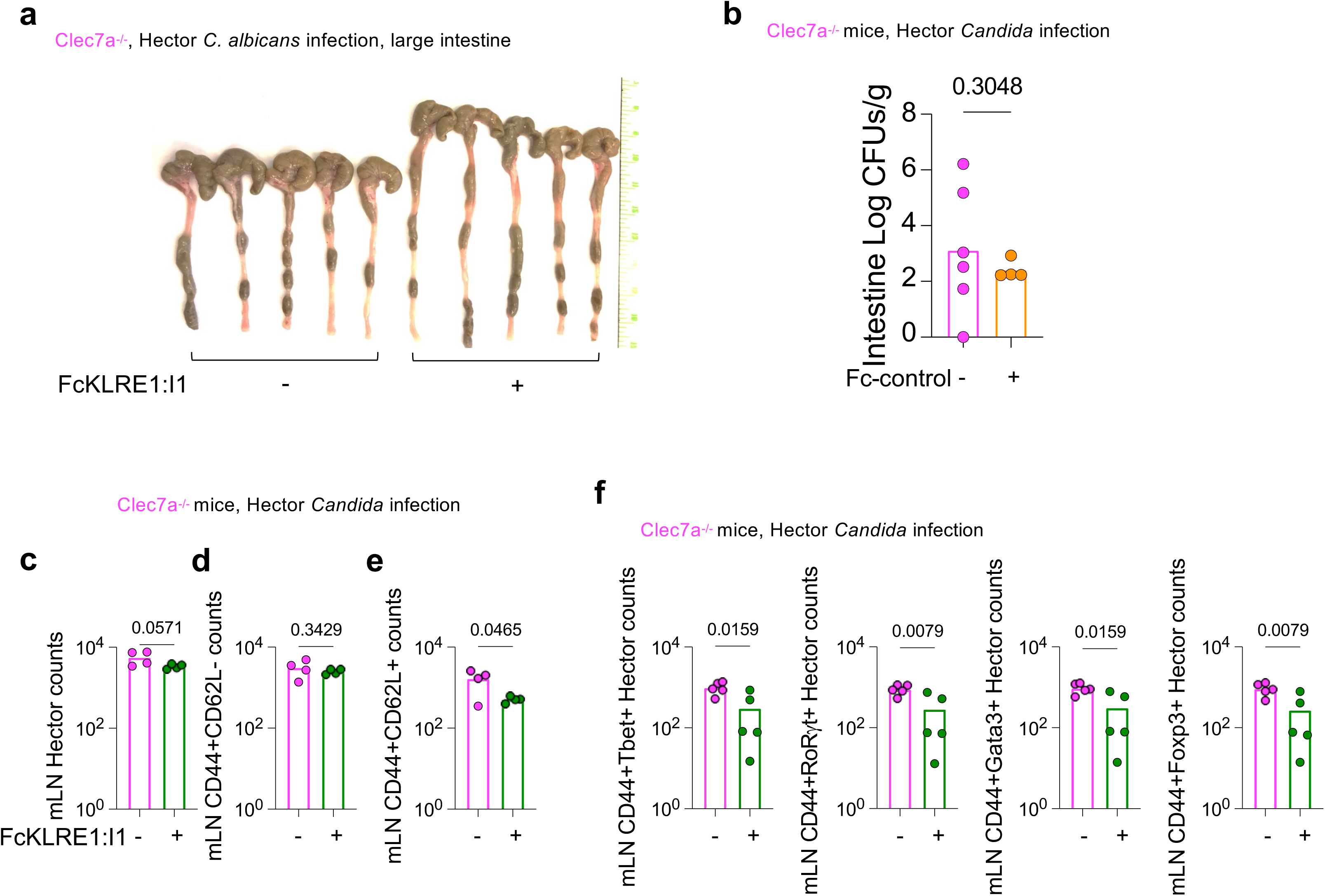
(**a**) Large intestines imaged at day 3 post infection from Clec7a^-/-^ mice with/out 7.5 ug/ml of FcKLRE1-I1 i.p given at day-1, and next day i.v infection with 1-1.5×10^5^ *C. albicans* SC5314 yeasts and a second dose of 7.5 ug/ml of FcKLRE1-I1 i.p given on day 1. (**b**) Intestinal fungal burden in Clec7a^-/-^ mice 3 days post-*C. albicans* infection following adoptive transfer of 1×10^6^ naïve Hector CD4^+^ T cells i.v. Mice received 7.5 µg/ml of unrelated Fc-control i.p at day-1 (one day before infection), followed the next day by i.v infection with 1×10^5^ *C. albicans* SC5314 yeasts, and a second 7.5 µg/ml i.p dose of the Fc-control treatment. mLN flow cytometric analysis of number of live (**c**) *C. albicans*-specific Hector CD4⁺ T cells; (**d**) effector memory CD44^+^CD62L^-^; (**e**) central memory CD44^+^CD62L^+^ and (**f**) numbers of antigen-specific CD44^+^Tbet^+^, CD44^+^RORgt^+^, CD44^+^GATA3^+^, and CD44^+^Foxp3^+^ cells. Each point represents one mouse; bars indicate mean. Data shown are representative of three independent experiments. Statistical analysis was performed using Student’s *t*-test with p ≤ 0.05 considered significant.

## Notes

### Competing Interest Statement

The authors have declared no competing interest.

